# Automated Protein Affinity Optimization using a 1D-CNN Deep Learning Model

**DOI:** 10.1101/2023.04.12.536512

**Authors:** J.Liam McWhirter, Abhishek Mukhopadhyay, Patrick Farber, Greg Lakatos, Surjit Dixit

## Abstract

Functional biologics design is a multi-objective optimization problem often with competing design objectives. We report on a novel deep learning based protein sequence prediction framework, ZymeSwapNet, that can be customized to handle a wide range of quantifiable design objectives, a current limitation of traditional protein design methods. We train a simple convolutional neural network (1D-CNN) on nonredundant curated protein crystal structures, using a set of geometric and topological features that describes a local protein environment, to predict the likelihood of each amino acid type for residue sites in the design region. While the model can be directly used to rank templates derived from mutagenesis campaigns, we extend the scope by developing a sequence/mutation generator that optimizes the desired multivariate distribution using a Monte-Carlo sampling. Using a case study – the design of a stable heterodimeric Fc (HetFc) antibody domain – we show that we can further include a Metropolis criterion to bias the sampling to enhance features such as the heterodimeric binding specificity, in addition to original sampling objective of enhancing stability. We demonstrate that ZymeSwapNet can generate stable HetFc designs, within minutes that had taken several rounds of rational structure and physical force-field based modeling attempts.

## 1. Introduction

The design of novel proteins has been traditionally a challenging and time-consuming problem, requiring searches over a huge space of structural and chemical changes. Such engineering efforts require multiple rounds of rational design, aided by traditional computational algorithms such as Dead-End Elimination [1], followed by experimental screening of mutations/designs. In rational design, protein engineers are also guided by chemical rules such as hydrogen bonds and electrostatic interactions which describe interactions limited to pairs and triplets of amino acid residues [2-3]. These rules are incomplete, unable to predict many body interactions between residues in a large group or network of residues. The protein optimization process is simplified by starting with a known folded protein structure and as such is a subset of the inverse-protein folding problem [4-24]: combinations of amino acid swaps are applied at different sequence positions on a fixed [17] or approximately fixed backbone [22], in the search for mutations that optimize a biological function, for example stronger binding affinity to a specified target, while maintaining stability. Even if a protein backbone is constrained, the space of possible designs is still enormous and very difficult to conceive how amino acid swaps at different residue sites/sequence positions will couple together in order to satisfy optimization objectives.

The optimization of functional proteins is a problem with competing objectives [19-24]. An example is the design of asymmetric bispecific antibodies [24, 25-30] relative to the symmetric chain composition of naturally occurring antibodies. Here the heterodimerization of two different Fc chains is a requirement for the formation of heavy chain heterodimers; the critical objective is to promote the formation of the Fc heterodimer (HetFc) while suppressing the formation of the two corresponding Fc homodimers [25, 28-30]. This is often referred to as multi-state design optimization [23]. In the traditional rational design process, achieving the first objective say achieving specific heterodimeric pairing can came at the cost of not satisfying a second objective, e.g. maintaining the stability of the Fc heterodimer. The ultimate satisfaction of both these objectives required several design rounds with a pool of trial mutations involving multiple residue sites (around 10 residue sites translating to ∼ N^10^ possible mutations where N < 20 is the number of physically reasonable amino-acid swaps per site).

Traditional computational protein design packages have two essential ingredients: a simulation / optimization method to sample/generate different protein backbone and side-chain conformations and a force-field / energy function for ranking the fitness of the generated protein structures. Often the protein backbone is assumed to be fixed and side-chain conformations are sampled from a library of observed crystal structures referred to as the rotamer library [12, 15]. Automated protein sequence design (APSD) has the additional ingredient where sequence identity is co-optimized along with structure. Besides the structure sampling algorithm, traditional APSD methods are limited by the quality of the energy function, its computationally efficiency and capacity to recover natural amino acid abundances in order to maintain the biological relevance of predicted protein sequences [14, 16, 17].

Building upon the work of a number research groups, the longstanding question of how to fold a protein given knowledge of only the protein’s sequence was recently “solved” by Google DeepMind’s Alpha-Fold [4-7]. Not surprisingly, there has also been significant research activity devoted to leveraging deep learning (DL) to solve the inverse-protein folding problem: neural network architectures have been trained on the vast wild-type sequence and structural data found in the protein databank (PDB) to predict protein sequence [31-40] given a fixed protein backbone conformation. APSD simulation methods can capitalize on these DL advances, relying on trained neural network models to significantly reduce the size of the sequence search space to the most-probable, potentially stabilizing mutations.

Early deep neural network (DNN) models for sequence design were simple, fully-connected neural network models with input features depending on only backbone geometry; these models relied on choosing a set of pre-engineered input features [31, 32]. More recently three-dimensional convolution neural networks (3D-CNN) [35, 38] and graph neural network [36, 40] architectures have been developed that have the capacity to learn the higher-order hierarchical relationships between input features, thereby reducing the need for feature pre-engineering. These models have been evaluated and compared based on their ability to recover the protein sequences given only the backbone structure as input. These models typically achieve wild-type sequence recovery rates of about 45 to 55 percent. For naturally occurring protein backbones, there is a distribution of sequences that fold into any given target structure, so it is not clear whether having a higher wild-type sequence recovery rate is especially meaningful above a certain threshold. Therefore, interest has shifted towards the generation of sequence distributions, that is, sequence diversity [38-40]. Of particular note is the recent work out of Stanford University by Anand et. al. [39]. Their 3D-CNN model was trained given all backbone atoms and side-chain atoms of residues neighbouring a target residue site within a fixed field of view; the side-chain atoms of the target site, pertaining to the target site were masked out, using the trained neural network to predict residue type and geometry of side-chain conformations at the target site. Sequence designs were then ranked via a heuristic energy function defined from the probabilities output by their neural networks [39]. Similarly, Ingraham et al from MIT [40] developed a graph transformer model that treats the sequence design as a machine translation from an input structure (represented as a graph) to a sequence: their model predicts amino acid identities sequentially using an autoregressive decoder given earlier decoded amino acid identities along the protein chain, proceeding one sequence position to the next in one decoding step, starting from one end of a chain and finishing at the other.

In this article, we report on the construction of a simple 1D-CNN model (ZymeSwapNet), trained to predict the amino-acid identity at a target residue site (Sections 2 and 3). The construction of ZymeSwapNet was inspired by work on the “Shared Residue Pair” (SRP) network [33, 37] Like SRP, ZymeSwapNet consists of two training stages and is trained given either backbone dependent or both backbone and side-chain dependent (Section 2) input features. The features are a coarse-grained representation of the protein where the side-chain dependent features are the amino-acid identities of the side-chains and exclude the side-chain atomic conformations.

Although ZymeSwapNet is trained to recover wild-type amino-acid identities (Section 3), we test whether heuristic energy metrics can be used to accurately score mutations, ranking these based on their ability to change protein stability or affinity with respect to wild-type (Sections 4.2 and 5.2). For affinity we compare ZymeSwapNet derived affinity changes to experimental protein-protein binding affinity measurements for a subset of mutations listed in the SKEMPI dataset [41]. Having established that these scores are sufficiently accurate and defined binding specificity metrics pertinent to HetFc optimization (Sections 4.3 and 5.3), we then construct a APSD simulation method that, for specified target region, generates novel mutations using a Monte-Carlo sampling algorithm (Sections 4.4, 5.4, 5.5). Residues in this target region are not required to be contiguous in protein sequence position, and the target region may span across multiple chains. This stochastic algorithm cycles across target residue sites: new amino acid identities at the target sites are assigned based on draws from the conditional probability distribution represented by ZymeSwapNet; as a result, the amino-acid sequence of the target region fluctuates during a simulation. We make the working assumption that upon convergence the sequence ensemble, generated by the recursively sampling from ZymeSwapNet across the target region, will be biased towards stabilizing sequences. To meet additional objectives, for example enhanced protein binding affinity or specificity, the sampling algorithm can be expanded to include a Metropolis criterion involving the additional protein property to enhance. In particular, we demonstrate the power of our approach by applying it to the selective design of an antibody Fc heterodimer (HetFc). We show (Section 5.5) that our in-silico approach can reproduce selective HetFc designs, such as “Knobs-into-Holes” (KiH) designs [25], within minutes that would otherwise take several rounds of traditional protein design and experimental verification.

## 2. Datasets and Features

### 2.1 PDB Structure Dataset

We selected two datasets from the Dunbrack Lab’s “Pisces” collection of culled PDB sequences [42]. The first dataset had chains (one chain identified per line) with a sequence similarity less than 90% (“pc90”) while the other had a similarity less than 30% (“pc30”). Both datasets had structures with a resolution less than 2.0 Angstroms, and a maximum R-factor of 0.25. A given PDB ID may have one or more chains occurring in a culled PDB dataset such that the number of unique chains in a dataset is greater than the corresponding number of unique PDB identities. We rejected structures if the following criterion were true: 1) structure was a protein-DNA or protein-RNA complex; 2) structure contained a D-amino acid; 3) structure contained residues with types ‘SEC’ (Selenocysteine), ‘ASX’ (ambiguous identity, either Asparagine or Aspartic acid), ‘GLX’ (ambiguous identity, either Glutamine or Glutamic acid), and ‘UNK’ (identity unknown); 4) the chain had less than 50 residues or more than 600. Hetatoms in a structure were ignored. The crystal structures were not optimized, that is, energy minimization was not performed in order to avoid any bias in the training of ZymeSwapNet.

### 2.2 Features Dataset Curation

For a given dataset, we created a corresponding ‘features dataset’. Each data array in a feature dataset is associated with a target residue site on a chain listed in the dataset. For each target site, we identified the K ‘closest’ neighbouring residue sites, based on C_*α*_ to C_*α*_ pair-residue distances. A neighbouring residue site may or may not be a target site associated with another separate data array. During validation, we set K to either 5, 10, 15, 20, 25, or 30, meaning that for K = 5 we defined a ‘cluster’ of 5 residues centred about a particular target site. A features dataset was then created from either the asymmetric unit ‘AU’ or the biological unit ‘BU’ crystal structures.

We grouped features into those that pertain to a single residue site (‘single-residue features’), and those that pertain to a target residue site and one of its neighbouring residue sites (‘pair-residue features’). When calculating features, we ignored side-chain atoms completely and treated only the protein backbone. A data array is related to a ‘star graph’ with the following feature group types:

1. Single residue features pertaining to the target residue site (set *S*_0_);
2. Single residue features pertaining to a neighbouring residue site (set *S*_*i*≠0_), ie, one of the ‘K’ nearest residue sites to the target residue site;
3. Pair residue features (set *P*_0*i*_) that depend on the geometry of the backbone segments of the target residue site and a neighbouring residue site.

### 2.3 Single-Residue Features

The set of all single-residue features pertaining to any given target site (Table 1) is denoted as 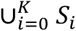 where *K* refers to the number of nearest-neighbours to target site ‘0’. There are 8 single-residue backbone features within a given *S*_*i*_: cosine and sine of 3 backbone torsion angles, secondary structure assignment, and solvent exposure. The secondary structure, determined using STRIDE [43], is a categorical variable; prior to training a deep learning model, we “one-hot-encoded” this feature. The STRIDE determination uses hydrogen bond patterns via an empirical energy function and backbone torsion angles. Solvent exposure was evaluated using DAlphaBall [44] given just the backbone atoms of a protein structure.

**TABLE 1:**
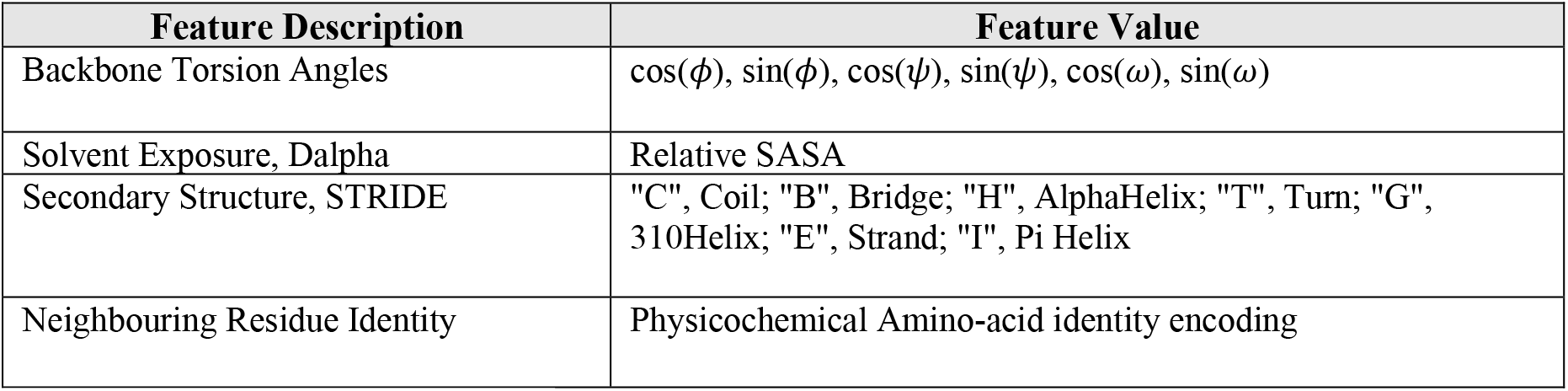
Single-Residue Features.

As a single-residue feature pertaining to a ‘neighbouring’ residue site, *S*_*i*≠0_, we also optionally included the neighbouring residue’s amino-acid identity. This amino-acid identity label (one-letter-code) could be one-hot-encoded into a 20-element long vector. However, applying this encoding creates a very sparse features data-array, which is memory expensive. Alternatively, we chose a mapping generated by a “symmetric neural network” used as an encoder-decoder. Once trained, this encoder subnetwork learns to represent a 7-D physicochemical properties vector as a 2-D vector; specifically, this encoder-decoder learns how to best regenerate an input 7-D vector given a training set of twenty 7-D physicochemical vectors, where each one of these twenty vectors corresponds to a particular amino-acid identity [45].

### 2.4 Pair-Residue Features

Pair residue features, denoted as *P*_0*i*_, *i* ∈ (1, *K*}, describe the positions of the neighbouring residue sites relative to the target site as well as the orientation of these neighbours. For any given K-neighbour cluster, we defined a reference configuration for the backbone segment of its target residue site, then found the transformation to move the target residue backbone segment from any starting configuration to this reference. The rigid K-neighbour cluster was then translated and oriented using this transformation. The reference configuration for the target was the following (an arbitrary choice adopted from Ref [33]): C_*α*_ atom is located at the origin, (0, 0, 0); N-C_*α*_ bond lies along the x-axis such that the x-coordinate of the N atom is negative; C backbone atom lies in the z = 0 plane such that the y-coordinate of the C atom is positive. After applying this transformation to the entire cluster, we calculated the K sets of pair residue features. The features within a given set *P*_0*i*_ are

1. Target C_*α*_ to neighbour C_*α*_ distance,
2. Unit vector from target C_*α*_ to neighbour C_*α*_ (x, y, z components of this vector),
3. Unit vector from neighbour C_*α*_ to neighbour N (x, y, z components of this vector), and
4. Unit vector from neighbour C_*α*_ to neighbour C (x, y, z components of this vector).

Features 1 and 2 give the position of a neighbouring residue site *i* relative to the target residue site ‘0’. Features 3 and 4 give the orientation of a neighbouring site *i* to the target site ‘0’. There are then 10 pair residue features for every possible target residue site to neighbouring residue site pair.

## 3 Neural Network Model

Given a set of input features associated with a target residue site, we constructed a neural network model, ZymeSwapNet, that maps these features to a classification/label of the amino-acid identity at this site. A data array of features, *X*_0_, associated with a given target site ′0′ and fed into ZymeSwapNet, has single-residue features *S*_*i*_, where *i* ∈ {0, *K*}, and pair-residue features *P*_0*j*_, where *j* ∈ (1, *K*}. Each node of the output network layer generates the probability of one of the 20 different (one-letter-code) amino acid identities possible via a “soft-max” output activation layer. Specifically, ZymeSwapNet outputs a vector of length 20 where a given element represents the conditional probability *P*(*Y* | *X*_0_) = *f*(*X*_0_; ***w, b***) of the “*Y*” amino-acid identity given the input features *X*_0_. Here ***w, b*** are the internal parameters determined by training.

### 3.1 1D Convolution Model

“ZymeSwapNet” (Figure 1) is a simple one-dimensional convolution neural network (1D-CNN). To train the 1D-CNN ZymeSwapNet, the input data example *X*_0_ is shaped into a 2D data-array:

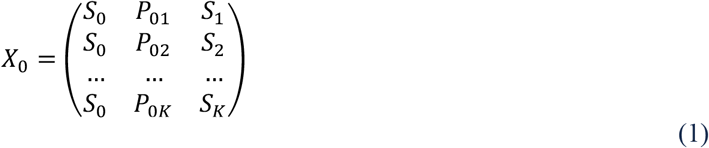

**FIGURE 1.**
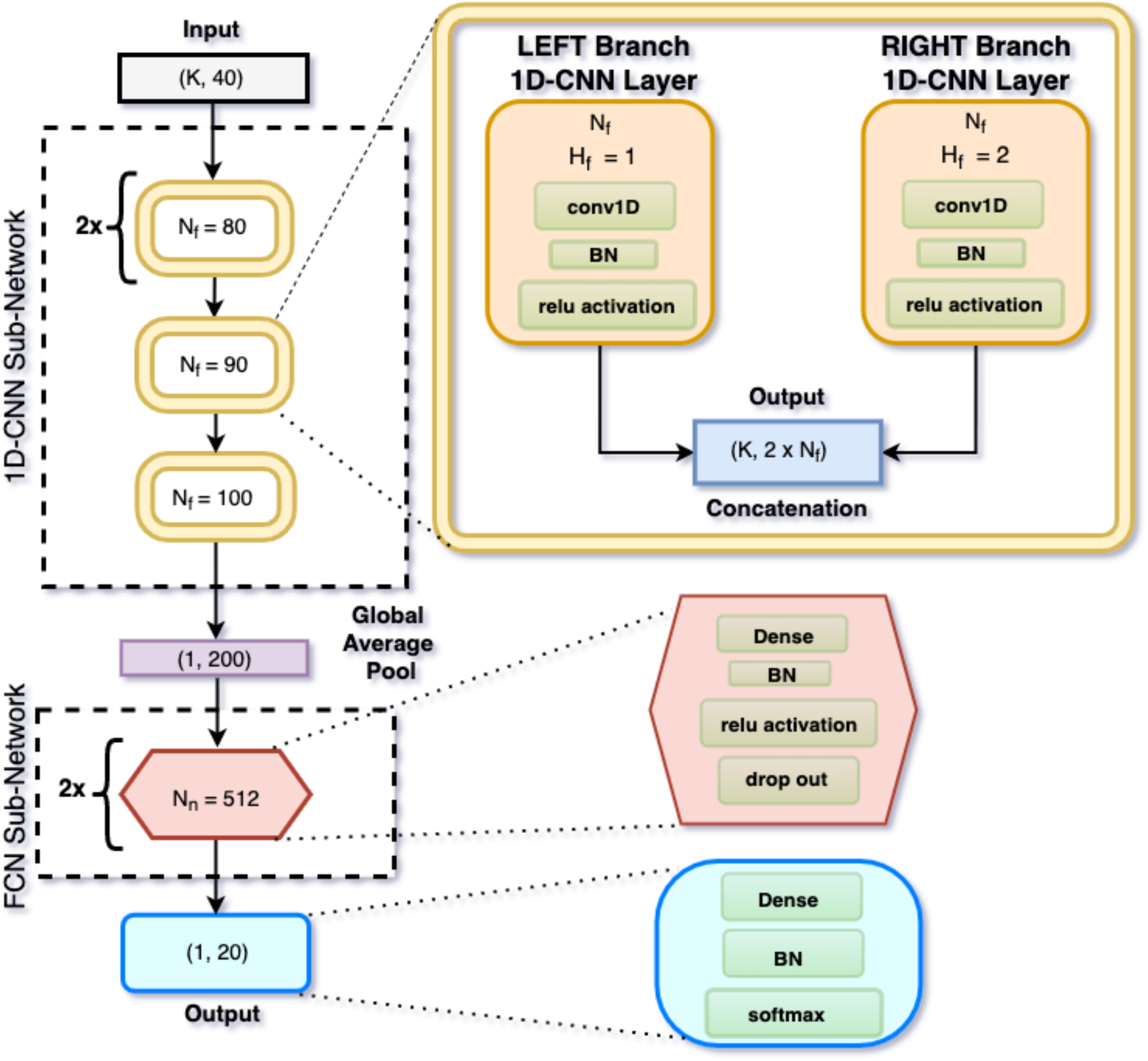
ZymeSwapNet model architecture: First, a 1D-CNN subnetwork consisting of four sequential levels, each with two parallel 1D convolution neural network (“conv1D”) layers, one layer in the left branch and another in the right branch; second, a FCN subnetwork consisting of two sequential levels, each with a single fully-connected neural network (“Dense”) layer. Dropout applied within the FCN sub-network is 20%. Each level has its own unique set of parameters to fit. For BB features, the shape of the input data array is (K, 40) where K is the number of residues in a neighbourhood surrounding the target residue site. The number and height of the filters in a 1D-CNN layer are respectively *N*_0_ and *H*_0_. The number of hidden activation units/nodes per FCN layer is *N*_*i*_. Batch Normalization (BN) was applied prior to any activation where the latter were either ReLu or softmax functions. The final softmax function outputs the probabilities at the target residue site of the 20 possible amino-acid identities.

Athough there is redundancy (*S*_0_ is repeated in the data-array), this data augmentation makes *X*_0_ amendable to 1D-CNN operations. ZymeSwapNet consists of two stages (Figure 1) where the first-stage is a simple one-dimensional convolution (1D-CNN) sub-network. This sub-network consists of four sequential levels with each level containing two parallel 1D-CNN layers (‘Left’ and ‘Right’ layers). At the end of a given level, after the activation stage of a convolution, the outputs of the two parallel, coincident 1D-CNN layers are concatenated laterally. Each 1D-CNN layer is characterized by the number of filters *N*_*f*_ in that layer, as well as the filter shape (Height, Width) = (*H*_*f*_, length(*S*_0_*P*_0*j*_*S*_*j*_)) where *S*_0_*P*_0*j*_*S*_*j*_ = *S*_0_ ∪ *P*_0*j*_ ∪ *S*_*j*_. For the ‘left’ and ‘right’ layers, the heights are *H*_*f*_ = 1 and *H*_*f*_ = 2 respectively. For both filter types, *H*_*f*_ = 1 and *H*_*f*_ = 2, the stride is size one. When striding, the filters run down the ‘height’ axis of a data array, applying a local linear transformation followed by a non-linear activation function at each stride step. If the shapes of the outputs from the two parallel, coincident 1D-CNN layers (one with *H*_*f*_ = 1 and the other with *H*_*f*_ = 2) are both ((*K* = 20), 80), then the lateral concatenation yields a data-array with shape ((*K* = 20), 160); this output then serves as input for the next two parallel, coincident 1D-CNN layers (again a ‘left’ and a ‘right’ layer) deeper into the network. Upon training, ZymeSwapNet, like other deep networks, should learn a hierarchy of progressively more complex features, where at each level in the hierarchy, features of two different resolutions / scope (*H*_0_ = 1 and *H*_0_ = 2) are “mixed”. Having *H*_0_ = 2 provides the network with a mechanism to learn/extract complex features involving the coupling of more than just 2 residues (ie many-residue interactions), that might prove advantageous for predicting the identity of the target residue. Going deeper into ZymeSwapNet, *N*_*f*_ per 1D-CNN layer increases, starting from the 3^rd^ level of the 1D-CNN sub-network *N*_*f*_ proceeds as 80 → 80 → 90 → 100. A 1D-Global Average pooling operation/layer is applied to the final concatenated data-array, collapsing this data-array along its height axis. This pooled data-array is fed into the second-stage fully-connected (FCN) sub-network which consists of 2 “Dense” layers each with *N*_*n*_ = 512 hidden activation units/nodes. All activations, with the exception of the output softmax layer, were ReLU functions, although we also experimented much later with eLU activations. Dropout regularization with a probability of 20% was applied only to the hidden activation layers of the FCN sub-network. Batch Normalization (BN) layers were applied immediately prior to any activation layer.

### 3.2 Model Training

ZymeSwapNet was trained given either just the backbone features (BB) or backbone features as well as the amino-acid identities of residues neighbouring a target residue site (BB + SC). When including “side-chain” (SC) features, the single-residue features set for a given neighbouring residue is simply appended to include the physicochemical encoding. In our notation going forward, (***X***_*t*_) will represent all backbone features associated with a target residue site denoted *t*. In addition, ***Y***_*neigh*(*t*)_ will represent the amino-acid identities of all K residues neighbouring a target residue site *t* ; these are the side-chain (SC) features. Therefore, a BB trained DNN model and a BB+SC trained DNN model represent the conditional probabilities *P*(*Y*_*t*_ | ***X***_*t*_) and *P*(*Y*_*t*_ | ***X***_*t*_, ***Y***_*neigh*(*t*)_) respectively of the amino-acid identity *Y*_*t*_ at the target site.

ZymeSwapNet internal parameters ***w, b*** were fitted by minimizing a Loss function (“categorical cross-entropy”) given the features dataset. The minimizations were performed using Stochastic Gradient Descent (SGD with ADAM) where one iteration of the descent acts on a mini-batch of training data with 128 input data arrays / objects / examples. For a few minimizations, the mini-batch size was increased to 32,768 data arrays; however, this made the accuracy of the amino-acid identity predictions about 4 to 5 percent worse. During a training run, SGD was performed for 100 epochs across the training data. Once 90 epochs were reached, we decreased the learning rate, effectively dropping the system into a local minimum of the Loss function. This drop is clearly seen in the plots of the Loss function and the accuracy as a function of epoch (Figure S1).

Based on PDB id, we used a 4:1 split to divide the features dataset into a development set and a hold-out set, then performed 5-fold cross validation on the development data, optimizing the value of K. In Table S1, we report 5-fold validation accuracies, and the prediction accuracies on the hold-out set with ZymeSwapNet retrained on the entire development set. For the Top-N accuracy, a successful prediction is counted as a true positive if any one of the Top-N most probable amino-acid identities predicted at the target residue site matches the actual label of that site in the PDB structure. Table S1 shows the Top-1, -2, -3, and -4 accuracies. For the (AU, pc30) dataset, the number of development and hold-out data arrays were about 1,200,000 and 250,000 respectively; the Top-1 Accuracy on the hold-out set after training ZymeSwapNet on the development set was 0.444 and 0.507 given BB and BB + SC features, respectively. For the (AU, pc90) dataset, the number of development and hold-out data arrays were about 1,800,000 and 350,000 respectively; the Top-1 Accuracy on the hold-out set was 0.477 and 0.533 given BB and BB+SC features, respectively. For both pc30 and pc90, the performance of the model was independent of whether we extracted an AU or a BU structure. All the results reported in following sections were derived from the (AU, pc90) ZymeSwapNet model.

## 4 Methods

### 4.1 Scoring Mutations using ZymeSwapNet

The 20 amino-acid probabilities output by ZymeSwapNet can be converted into a heuristic energy, by taking the −ln(·) of these probabilities [32, 33]. We propose three DNN-based scoring / energy functions. These energies can rank the stabilities and affinities of different sequence designs. The neural network is trained to predict the amino-acid identity *Y*_*t*_ at a given target residue site *t* given backbone features ***X***_*t*_ and the amino-acid identities of the nearest-neighbouring residues to the target site, ***Y***_*neigh*(*t*)_. These input features are a coarse-grained representation of the protein. However, this network outputs a conditional probability for a single site, *P*(*Y*_*t*_ | ***X***_*t*_, ***Y***_*neigh*(*t*)_), and not a joint probability *P*(***Y***_2_ | ***X***_2_, ***Y***_*neigh*(2)_) for a given target region, *T*, where 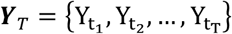 and *t* ∈ {*t*_1_, *t*_2_, …, *t*_2_}. A neighbour of any given target residue site, *τ* ≠ *t, τ* ∈ *neigh*_*t*_, will be either itself a target site in the target region or a background/environment residue site. Hypothetically if we had the joint distribution available, we could insert a sequence of interest into the -ln(*) of this distribution and then use such a scoring method to rank all sequences of interest with respect to stability. To reconstruct a neural network so that it outputs a joint probability for a sequence/set of target sites involves re-architecting the existing network as a component/sub-network of a larger *Conditional Random Field* (CRF) network [46-48]. Switching to a CRF, training examples go from being target sites to target regions. CRFs were not investigated in our work.

Despite not having absolute scores associated with some hypothetical joint distribution, we can still pose an approximate expression for the difference in the scores/energies between any two sequences for a given target region (for example, a final mutant sequence and a starting wild-type sequence). Outside the target region, the background/environment residue sites have their amino-acid identities, ***Y***_*bkg*_, kept fixed. Going from an initial sequence 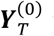 to a final sequence 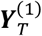, we evaluate the change in stability due to this mutation by sequentially applying each amino-acid swap in the mutation (ie the “mutation swaps”) to the target region, summing the incremental changes from each individually applied swap:

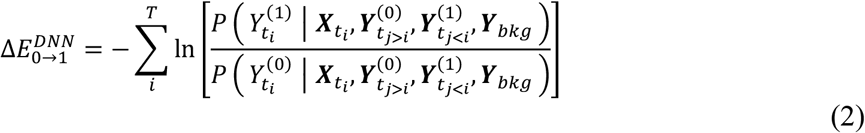

A first approximation replaces the numerator and denominator of the ln() argument above with the DNN representation of the conditional probability distribution, where in addition to the backbone (BB) features, ***X***_*t*_, the DNN distribution depends on the amino-acid identities of the K nearest-neighbouring residues to the target site, *t*. After applying a swap to a target site, all of its neighbouring residue sites must have their input features data arrays updated to reflect the change in amino-acid identity at *t*: there are non-additive effects between swaps. Given the DNN approximation, with its fixed, finite value of K, the sum becomes dependent on the order that the swaps are applied; therefore, it is a rational decision to average over a number of randomly chosen permutations to the order of the applied swaps. Each permutation represents a “mutation path”. One can make an additional ad-hoc approximation: when applying a swap to a given target site, the amino-acid identities at all other residue sites are set to their wild-type identities in the DNN approximation. This “wild-type” approximation amounts to the working assumption that swaps are decoupled and have just an additive effect; this assumption is expressed by replacing 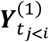 by 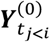. In short, when both backbone (BB) and side-chain (SC) features are input into the DNN model, the swaps can be treated as being either coupled or decoupled. With just backbone (BB) features, one swap is independent of any other swap, and so the total stability change, ie, the score relative to the starting sequence, will not depend on the order of the applied swaps. We will refer to the stability change due to mutation calculated by Equation (2), when the swaps are coupled or decoupled, as respectively the “PathAve_DNN” and “Decouple_DNN” stability change metrics.

The second two DNN scoring functions, which we will refer to as the “Local_DNN” and “Global_DNN” energies, are calculated given the instantaneous sequence state, such that the “absolute energy” is *defined* as

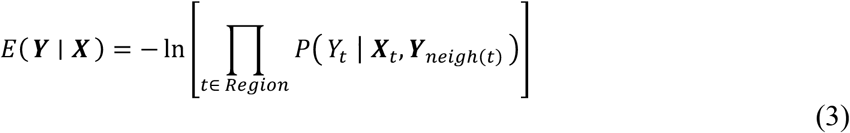

This “Region” can be restricted either to 1) a local region composed of only those residue sites corresponding with the mutation swap sequence positions (“Local_DNN” energy), or 2) the entire protein structure, for example, the entire complex, ligand, or receptor (“Global_DNN” energy). These are both heuristic energy functions: in general, a joint probability distribution of several variables does not factorize into the product of conditional probabilities in Equation (3); specifically, Equation (3) does not follow from chain-rule of probabilities. Although not conceptually as rational as the “PathAve_DNN” stability change metric, these two absolute DNN energies, as well as the stability and affinity metrics derived from them, are much faster to calculate. Protein-protein binding affinity changes due to mutation predicted via changes to the “Local_DNN” and “Global_DNN” affinity metrics correlate well with corresponding predictions made by the “PathAve_DNN” affinity change metric (Section 4.2 and Section 5.2, Figure 3).

### 4.2 Scoring Mutations: Affinity and Stability

Scoring remains a major challenge while modelling properties like affinity and stability using traditional molecular modelling and simulation methods. We have explored the utility of the DNN-based scores for predictions of affinity and stability changes, due to mutation from wild-type, and compared several different choices of stability metric for two experimental (SKEMPI and FAB) datasets. The affinity of a protein complex with a ligand, protein ‘A’, and a receptor, protein ‘B’, is

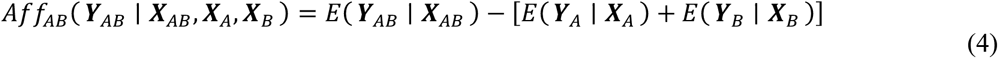

and the change in affinity due to mutation from a wild-type or some initial state is simply

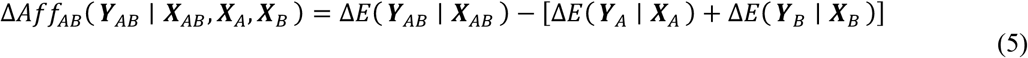

where the change in stability due to the mutation is Δ*E*(***Y***^*mut*^ | ***X***) = *E*(***Y***^*mut*^ | ***X***) − *E*(***Y***^*wt*^ | ***X***). One set of stability metrics are DNN-based. The other set are traditional, either physics-based or knowledge-based (we will refer to these collectively as “physical”), and require information about the conformations of side-chain atoms which we optimized using rotamer packing methods [12, 49] followed by short runs of deterministic minimizers such as steepest descent and L-BFGS to remove strong atomic clashes (our “structural repacking workflow”; Section S.2). Our physics-based energy function is the Amber99 force-field augmented with 1) a Generalized Born (GB-OBC) energy term to model the solvent polarization affect on Coulombic interactions between protein atoms [50], and 2) a simple interatomic pairwise Hydrophobic energy term that favours interactions between non-polar atom types [51]. We will refer to this function as the “Amber” energy. Our knowledge-based energy function was determined gathering statistics of atom-pair distances from a database of high-resolution, non-redundant protein structures (PDB dataset, Section 2.1). This energy function has two terms: a “Distance-Dependent Random Walk” reference potential term [52], and a “HClash” term that detects clashes between hydrogen atoms and any heavy atoms in the structure using a Lennard-Jones (LJ) like expression. We will refer to this function as the “DDRW” energy.

### 4.3 Scoring Mutations: Binding Specificity

We make binding specificity predictions based on ZymeSwapNet derived metrics, comparing the affinity of a Fc heterodimer, *A*_1_*B*_2_, to the affinities of two corresponding homodimers, *A*_1_*B*_1_ and *A*_2_*B*_2_. If a set of swaps *S*_1_ is applied to chain A and another set *S*_2_ is applied to chain B to form the heterodimer *A*_1_*B*_2_, then the two corresponding homodimers are formed respectively by applying *S*_1_ to both chains A and B (yielding homodimer *A*_1_*B*_1_) and applying *S*_2_ to both chains A and B (yielding homodimer *A*_2_*B*_2_). ZymeSwapNet was applied to the two chain Fc domain structure, AB, as well as to each individual chain, A and B, where each chain was isolated/unbound from the other. To derive a measure of the stability change of the bound species given a mutation (ie, a set of swaps), ZymeSwapNet probabilities associated with the complex AB were extracted, then the DNN-based energies were evaluated from these probabilities. Similarly, to derive the stability change of the unbound species given a mutation, the neural network probabilities and DNN-based energies associated with each individual chain, A and B, were used.

If a DNN-based affinity metric is qualitatively accurate, then for Fc designs/mutations that yield “pure” HetFc lab samples we would expect a larger drop in “energy” upon mutation going from the unbound to bound complex state for the Fc heterodimer than for the two corresponding Fc homo-dimers. In principle, this larger drop should be reflected by a more negative value of a binding specificity metric. We define such a metric using either the average (func() = ave()) or minimum (func() = min()) function:

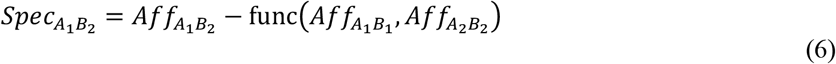

We define a “change” in the binding specificity metric due to mutation with respect to a starting wild-type sequence as

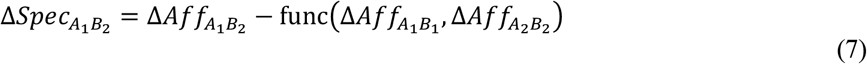

Given the expression for 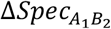 above, 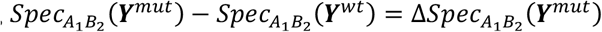 is true when either func() is the “Minimum” function and all chemical species (either complex, ligand, or receptor) share the same wild-type reference (either complex, ligand, or receptor, respectively), or when func() is the “Average” function. Furthermore, if the “absolute energy” is the “Global_DNN” energy, then since all three complexes, the heterodimer and two homodimers, share the same wild-type reference, the change in the binding specificity from wild-type due to mutation will depend only on the mutant and not the wild-type sequence: 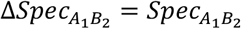.

### 4.4 Automated Binding Specificity Optimization

We can score designs but we also want to automatically generate them using ZymeSwapNet. In this section, we present a computational simulation method that, given a fixed protein backbone, generates sequence stable HetFc designs that favour the formation of a Fc heterodimer (HetFc) over the formation of two corresponding Fc homodimers.

#### 4.4.1 Simulation Method

For a given target region on the heterodimer Fc, *T*, the algorithm uses Gibbs Sampling to effectively sample the unknown joint probability, *P*(***Y***_2_ | ***X***_2_, ***Y***_*neigh*(2)_), of the amino-acid sequence of labels 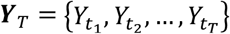 by successively sampling the associated known conditional probabilities, *P*(*Y*_*t*_ | ***X***_*t*_, ***Y***_*neigh*(*t*)_) where *t* ∈ {*t*_1_, *t*_2_, …, *t*_2_}. Prior to sampling, ZymeSwapNet is trained to predict the amino-acid identity *Y*_*t*_ at a given target residue site *t* given backbone features ***X***_*t*_ and the amino-acid identities of the nearest-neighbouring residues to the target site, ***Y***_*neigh*(*t*)_. A neighbour of any given target site will either be itself another target site in the target region, whose amino-acid identity/label fluctuates during sampling, or a background/environment residue site, whose amino-acid identity/label is kept fixed. Sampling from ZymeSwapNet ensures that we are drawing an amino-acid identity that is instantaneously likely and so, we assume, stabilizing for at least the Fc heterodimer. However, the swap is not immediately accepted because there is an additional objective besides stability to optimize, the binding specificity of the Fc heterodimer.

We want to generate designs where the binding of a Fc heterodimer (ie, *A*_1_*B*_2_) is preferred over the binding of the two corresponding Fc homodimers (ie, *A*_1_*B*_1_ and *A*_2_*B*_2_, refer to Section 4.3). To optimize (on average) this additional objective, we only accept an “attempted” swap (ie, change in amino-acid label at a given target site) conditionally based on a Metropolis criterion. The imposition of this criterion means that a specificity bias is explicitly “turned-on”. If the “change” in the specificity arising from an attempted swap, Δ*Spec* is negative, Δ*Spec* ≤ 0, then we accept the swap outright; otherwise, if Δ*Spec* > 0, then we accept the attempted swap only if a sampled uniform random number *R*, with value between 0 and 1, is less than exp(−Δ*Spec*/η). We have added an artificial temperature η to control the swap acceptance rate.

The Monte-Carlo (MC) simulation algorithm involves cycling through the target region of the Fc heterodimer. At the start of each cycle, the path to follow through the target residue sites of the target region is chosen at random. During a single cycle, the algorithm visits one target residue site at a time along this path until all sites have been visited. At each target site visit, a swap is attempted: first, the swap is drawn by sampling the probability distribution output by ZymeSwapNet, then this attempted swap is either accepted or rejected according to the Metropolis criterion discussed above. For sampling, the residue sites *t*_*i*_ of the target region are NOT restricted to being contiguous in sequence space. If a swap is accepted, then the amino-acid identity of the current target site is updated as well as the features vectors of any neighbouring residue sites, ie, those residue sites which have the current target site as one of their K nearest-neighbours. The latter update is necessary because the amino-acid identity of neighbouring sites have been chosen as features (BB+SC). Because our objective is to enhance the binding specificity towards the Fc heterodimer, *A*_1_*B*_2_, in order to calculate the “change” in the specificity, Δ*Spec* (*as defined* by the expression in Section 4.3), we also need to calculate the change in affinities of the two corresponding homodimers *A*_1_*B*_1_ and *A*_2_*B*_2_. Therefore, for given swap α → β we must also calculate the change in the DNN stability metric for each bound protein species (*A*_1_*B*_2_, *A*_1_*B*_1_, *A*_2_*B*_2_) as well as for each unbound species (*A*_1_, *A*_2_, *B*_1_, *B*_2_). Each of these seven species will require a distinct set of feature data arrays, updated accordingly should the attempted swap be accepted. Every target site *t*_*i*_ on the Fc heterodimer may or may not have a corresponding “target site” on any one of the remaining complementary bound and unbound protein species. The change in the stability metric due to an attempted swap for any given species, bound or unbound, from an initial amino-acid identity 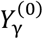 to a final identity 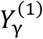 is simply defined as

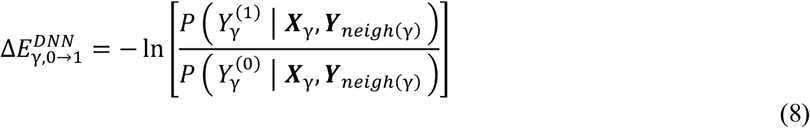

where γ denotes either the target site *t* on the heterodimer or a residue site associated with this target site on the remaining 6 protein species. This incremental change in stability due to a swap has the same expression as one of the terms in the evaluation of the “PathAve_DNN” stability change metric (Section 4.1).

#### 4.4.2 Simulation Procedure

A MC simulation run consisted of 2000 Gibbs cycles through an entire target region on the Fc heterodimer. Each run began by randomizing the amino-acid identities of each target residue site for all protein species involved: the bound complexes and the unbound ligands (chains A) and receptors (chains B). The first 500 cycles were discarded from the analysis; these were associated with a relaxation period to a “steady-state” where the DNN-based stability scores rapidly improve. Taking sequence “snapshots” every 10 cycles, we then clustered the resulting 150 *A*_1_*B*_2_ sequence designs (sequence snapshots from the simulation trajectory) using hierarchical clustering, generating dendrograms. Typically as the specificity bias/drive was increased, large clusters (ie, clusters containing more than 10 (an arbitrarily number) unique Fc heterodimer *A*_1_*B*_2_ sequence designs) became increasingly more selective for HetFc. This trend was reflected by a decreasing value of a binding specificity metric, 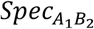: a lower value represents a more favourable tendency to form the Fc heterodimer over the two corresponding Fc homodimers. Sequence similarity scores, required in order to assign cluster identities, were calculated based on a “fine-grained” grouping of amino-acid identities (Section S.3, Table S3).

The artificial temperature was set to either η = 1.0, η = 0.5, or η = 0.25. The specificity bias in the MC was also either turned on (weight, W = 1) or off (weight, W = 0). When W = 0, the temperature had no effect because we chose to not add a temperature control to modify the ZymeSwapNet probabilities. Note having the binding specificity bias explicitly *turned off* did not mean that there could not be a selective drive to form a Fc heterodimer (HetFc): if we chose to optimize the sequence of a target region that straddles the heterodimer interface (with some residues on chain *A* and others on chain *B*), then sampling swaps from the conditional probabilities output by ZymeSwapNet acting on the HetFc will stabilize the HetFc likely by increasing the binding of the HetFc at some cost to the binding of the two corresponding Fc homodimers.

## 5 Results and Discussion

### 5.1 Model Test: Antibody Fc Domain Predictions

We applied the DL model to an antibody CH3 domain Fc interface in order to predict mutations that drive heterodimerization. The predictions are presented in Table 2. It is encouraging to see that many of the stabilizing swaps that were earlier (Ref 29) obtained using multiple rounds of rational engineering are to amino-acid identities that are more probable, as predicted by ZymeSwapNet, than the corresponding wild-type amino-acid identity.

**TABLE 2.** ZymeSwapNet Amino-acid Identity prediction for the protein structure, IgG1 Fc domain (3AVE). For a given row, pertaining to a swap/residue site, predictions are ordered from less to more probable, going from left to right. Amino-acid identities which are highlighted in red contribute to about 70 % of the total probability, ie, the sum of their probabilities is about 70 %. Mutant amino-acid identities taken from the protein engineered sequence designs [29] are highlighted in bold. Wild-type amino-acid identities (if they even appear in a list) are underlined.

| Swap Site | Design Swap | Top Amino Type DNN Prediction, BB | Top Amino Type DNN Prediction, BB + SC |
| --- | --- | --- | --- |
| **A350** | T350V | SER, <u>THR</u>, **ALA**, **VAL** | SER, ALA, <u>THR</u>, **VAL** |
| **A351** | L351Y | TRP, HID, **PHE**, **TYR** | HID, <u>LEU</u>, **PHE**, **TYR** |
| **A405** | F405A | GLN, **ALA**, **THR**, **SER** | **THR**, **ARG**, **ALA**, **SER** |
| **A407** | Y407V | TRP, <u>TYR</u>, **ALA**, **SER** | **ASP**, **SER**, <u>TYR</u>, **GLY**, **ALA** |
| **B350** | T350V | **SER**, **VAL**, <u>THR</u>, **ALA** | **CYS**, **ILE**, <u>THR</u>, **VAL** |
| **B366** | T366L | GLN, **GLY**, **MET**, **LEU** | **LEU**, **MET**, **GLY**, **GLN** |
| **B392** | K392L | **LEU**, **THR**, **ILE**, **VAL** | **ILE**, **SER**, **THR**, **LEU**, **ASN** |
| **B394** | T394W | **ALA**, **CYS**, **SER**, <u>THR</u> | **ALA**, **ASP**, <u>THR</u>, **SER** |

For the next test, an entropy is defined to describe the shape of a probability distribution of the 20 different amino-acid identities at any given residue site,, targeted for mutation:

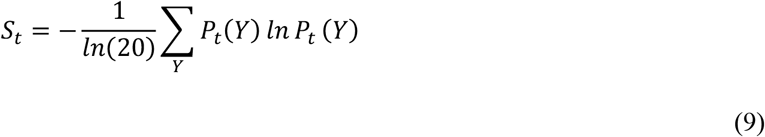

Here ∑_*y*_ *P*_*t*_(*Y*) = 1 such that *P*_*t*_(*Y*) = *P*(*Y* | ***X***_*t*_, ***Y***_*neigh*(*t*)_). As the entropy increases, becomes flatter in shape, ie, more uniform. If we assume that ZymeSwapNet accurately encodes information about general patterns found from the PDB, then a uniform distribution at a target residue site signifies that one choice of amino-acid identity is as good as any other. Conversely, as the entropy decreases, becomes more peaked over particular amino-acid identity, indicating a strong prediction of this identity. As an example of this scoring procedure, swapping a THR for a large “Knob”, TRP, at position 394 on chain B changes the entropy the most at a neighbouring residue site across the interface on chain A, at position 407. Figure 2 suggests that this B/T394W swap, in principle, could be stabilized by also adding a complementary swap to either a GLY or ALA at position A/407, which being small residues would represent the formation of a corresponding “Hole”. Introducing the B/T394W swap significantly reduced the entropy only at site A/407.

**FIGURE 2.**
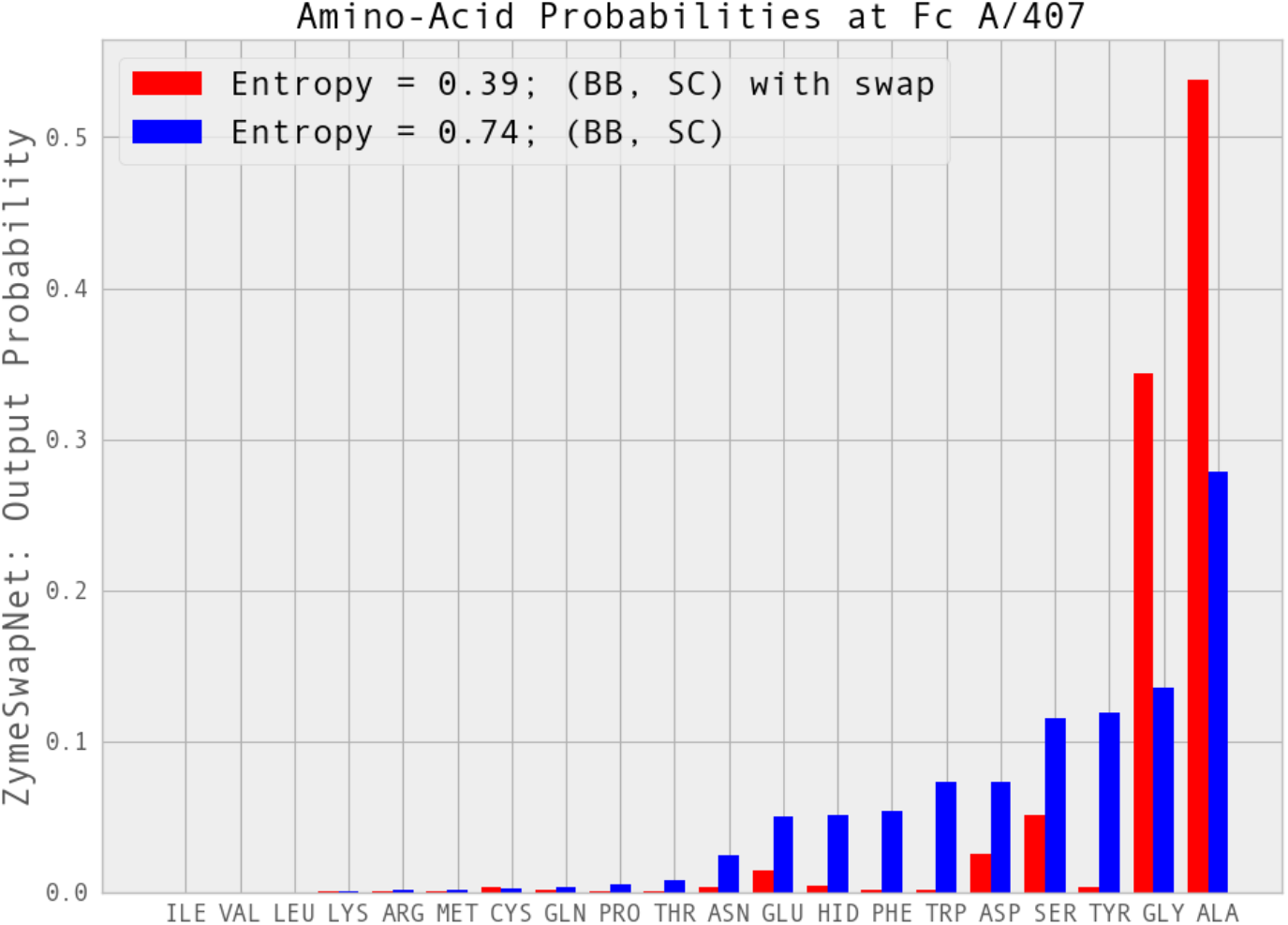
Amino-acid probabilities at site A/407 of IgG1 (3AVE) given swap B/T394W. Adding a “Knob”, then finding a “Hole”. Probabilities were generated from ZymeSwapNet trained given backbone (BB) and side-chain (SC) features. In the legend, entropy values at position/site A/407 are shown prior to and after the application of the B/T394W swap from THR to TRP at sequence position B/394.

### 5.2 Scoring Mutations: Affinity and Stability

The first experimental dataset consists of data for 3014 mutations, generated from the SKEMPI database of protein-protein binding affinity changes [41]. Comparing the predictions of the affinity changes, we see that the DNN-based and “physical” affinity metric changes due to mutation correlate well with each other, as well as with experimental measurements of ΔΔ*G*. Despite this success, at least for this dataset, the DNN-based affinity metric changes were not affected by the addition of coupling between separate mutation swaps (compare in Figure 3 the correlation coefficients involving the DNN-based affinity metrics). To create Figure 3, the “PathAve_DNN” stability change metric for any given mutation and protein species (complex, ligand or receptor) was calculated by averaging over 50 or 200 randomly chosen permutations (“mutation paths”) to the order of the applied swaps (Section 4.1 and 4.2). However, again, at least for our curation from SKEMPI dataset, correlations involving the “PathAve_DNN” affinity metric changes appeared largely independent of the number of these paths (50 or 200).

**FIGURE 3.**
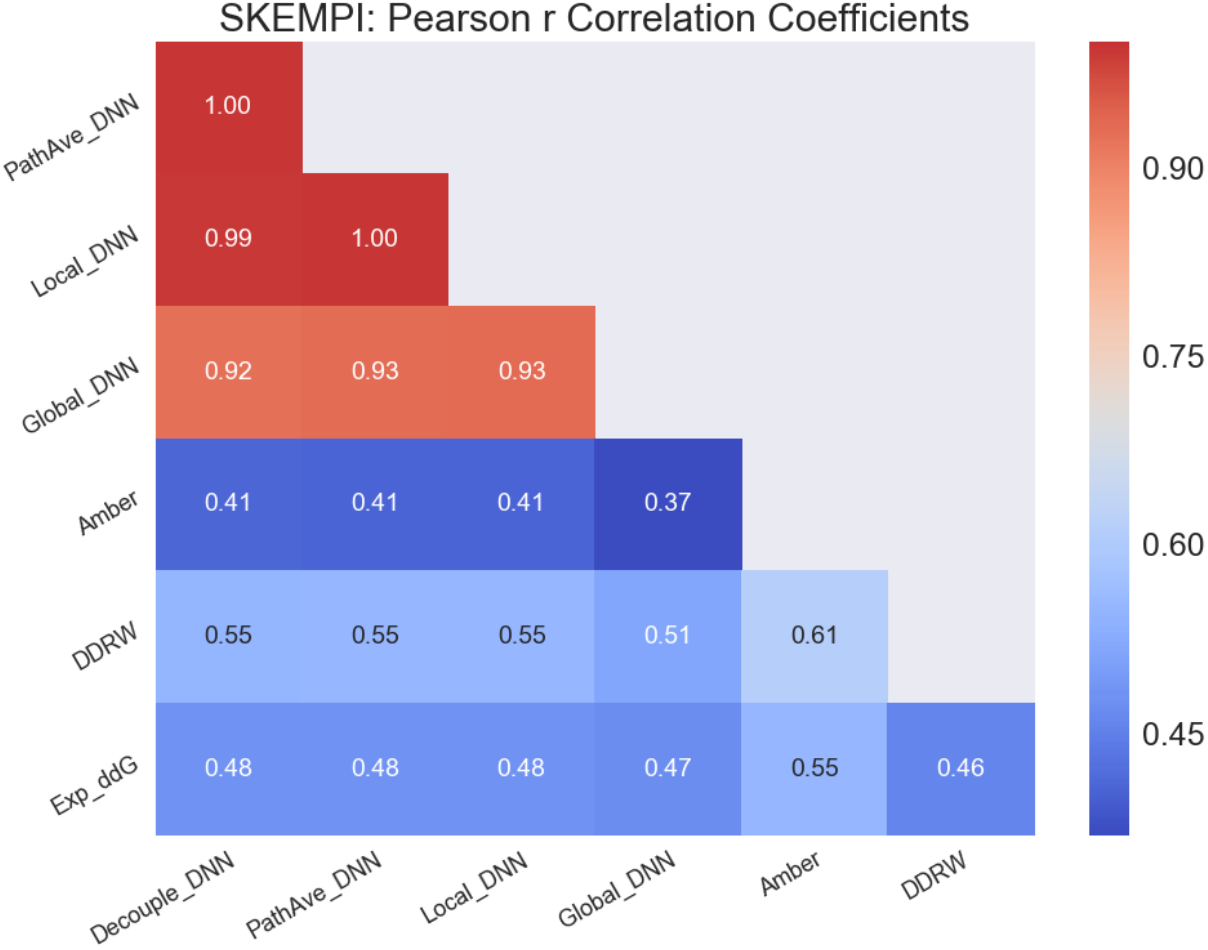
Correlations (Pearson r) between predictions (DNN-based, physics-based and knowledge-based metrics) and experimental measurements (SKEMPI dataset) of protein-protein binding affinity changes due to mutations from wild-type. All DNN-based metrics employed neighbouring amino-acid identities as features (ie, BB + SC, see Section 3). For the calculation of the “Decouple_DNN”, swaps within any given mutation were treated as being decoupled: here when calculating the swap contribution to the stability change, residues neighbouring the swap site had their amino-acid identities set to wild-type. Correlation coefficients dropped significantly when a ZymeSwapNet model trained only on BB features was used to calculate DNN-based metrics (not shown). Refer also to Figure S4.

The second experimental dataset consists of internal data for 390 mutations, curated from experimental measurements of melting temperature changes, Exp_dTm, due to mutations of a Fab antibody domain (PDB id, 1JPT). Unlike the mutations in the SKEMPI dataset, for the FAB dataset turning-on the coupling between mutation swaps significantly increased the correlations between the DNN-based stability metric changes and experimental measurements (Figure S5). Once again, the correlations to the “PathAve_DNN” stability change metric did not appear to depend on the chosen number of mutation paths to average over in the calculation of this metric. Nonetheless, as we will show in Section 5.3, when calculating the “PathAve_DNN” stability change metric, changing this number had a significant effect when calculating binding specificity metric changes due to mutations. Therefore, as a policy, it appears prudent to average over mutation paths when calculating a “PathAve_DNN” metric.

### 5.3 Scoring Mutations: Binding Specificity

Figures 4 and S6 show the 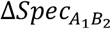 derived from the “PathAve_DNN” stability change metric. Backbone and side-chain dependent features (BB + SC) were input into ZymeSwapNet. We compared binding specificity metric changes observed when the coupling between swaps was disabled (ie, each swap sees their neighbouring residues as having wild-type amino-acid identities) and enabled. Mutations were taken from an internal “HetFc” stability dataset consisting of the experimental melting point measurements for ∼100 mutations applied to the IgG1 Fc domain (PDB id, 3AVE) that yielded a strong preferential formation of the Fc heterodimer. For Figures 4 and S6, the 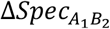 were calculated using respectively the “Average” and the “Minimum” specificity functions. When the swaps were coupled, the number of permutations to the order of the applied swaps (ie number of mutation paths) was increased gradually from 1 to 100. Qualitatively the plot appearance converged once this number exceeded around 50. Each point within Figures 4 and S6 corresponds to one particular HetFc design / mutation. The mutations were divided into three groups, each associated with a different experimental % purity (85 %, 90 %, and 95 % pure) of the Fc heterodimer.

**FIGURE 4.**
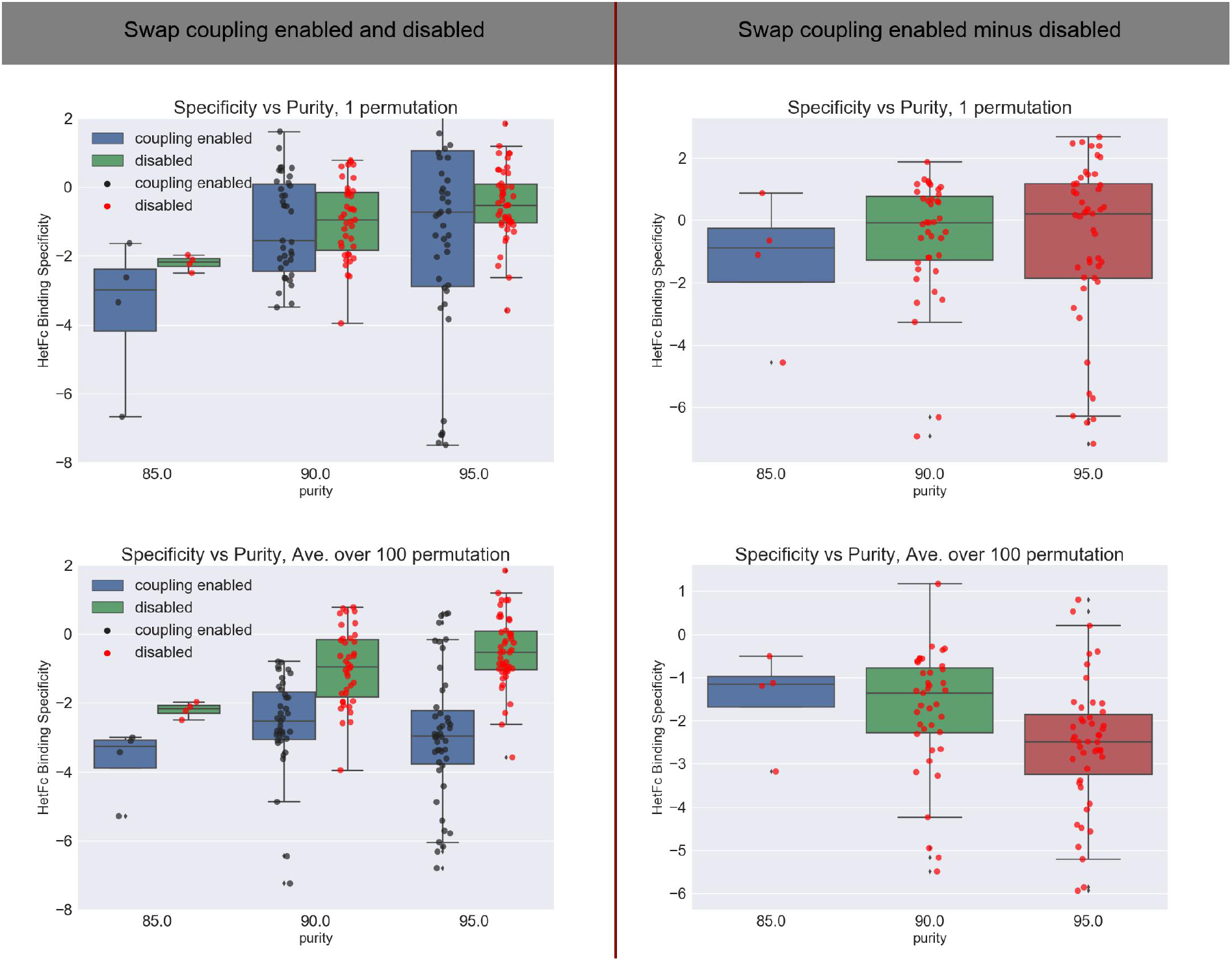
“Ave” Specificity metric changes due to mutation in the HetFc stability dataset. Comparison of binding specificity changes between runs with and without coupling (“non-additive” effects enabled and disabled respectively) between multiple swaps in a mutation. “PathAve_DNN” stability change metrics were calculated from an average over a single (first row) or 100 (second row) mutation paths. The “Delta calculation” (second column) for a mutation equals the binding specificity metric evaluated with coupling between the swaps minus the binding specificity metric evaluated without coupling between the swaps. Refer also to Figure S6.

All HetFc dataset mutations/designs yielded lab samples of very high purity (85 to 90 %) in the Fc heterodimer. Ideally, therefore, both of the two defined binding specificity metric changes (func = “Ave” for average, and func = “Min” for minimum) due to mutation should be negative in value for most of the Fc heterodimer mutations; however, this appeared only true for the func = “Ave” binding specificity metric (Figure S7). The “Delta calculation” in the second columns of Figures 4 and S6 shows that the inclusion of coupling between mutation swaps delivers a negative contribution to the change in the “Ave” and “Min” binding specificity metrics, driving the specificity values towards the expected, negative valued range. From this “Delta calculation”, there is an additional trend where as the sample purity goes up the non-additive contribution becomes a more negative number; however, this trend doesn’t appear as distinct for the “Local_DNN” metric or at all for the “Global_DNN” metric (not shown). What is evident from these plots is the benefit of averaging over mutation paths when trying to predict an expected qualitative value for 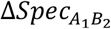 calculated using the “PathAve_DNN” stability change metric: including non-additive effects and averaging over swap permutations produces a more negative value for the “Ave” and “Min” 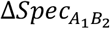.

For the IgG1 Fc domain (PDB id, 3AVE), critical KiH swaps consisted of a swap T394W on chain B and a pair of complementary swaps F405A and Y407V on chain A [29]. As desired, turning-off some of the critical KiHs swaps listed in [29] affected the “Ave” and “Min” 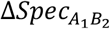 (Figure 5). If our DNN-based affinity metrics truly capture the physical effects of these critical swaps, then removing these swaps should result in an unfavourable increase in the value of a 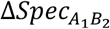. To turn-off or disrupt a critical KiH design, we simply removed the swaps involving A/405.PHE→ALA or B/405.PHE→ALA from a mutation/design and simultaneously removed B/394.THR→TRP or A/394.THR→TRP. Table S2 illustrates this edit for one example of a HetFc mutation/design from the HetFc stability dataset. As desired, turning off some KiH swaps increased the value in the “Ave” and “Min” 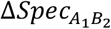.

**FIGURE 5.**
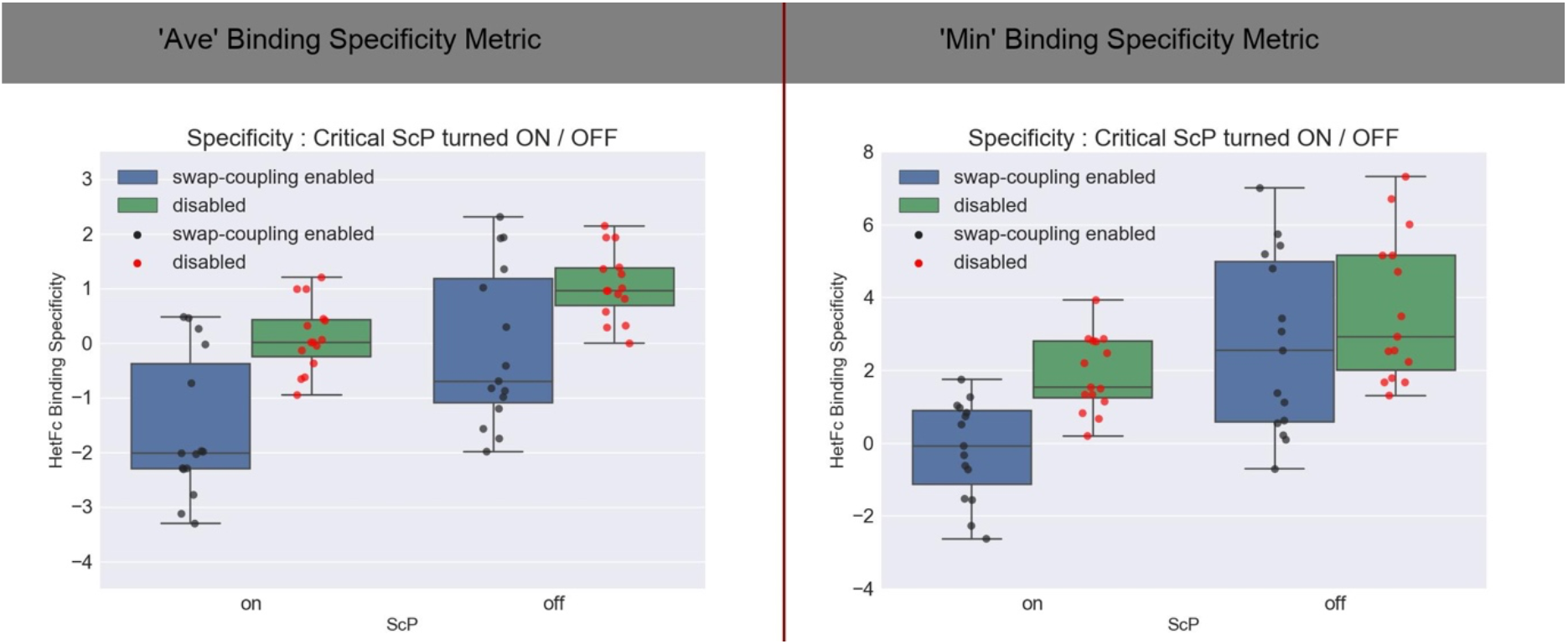
Effect of turning-off KiH swaps from HetFc mutation/design. Comparison of values of the changes to the “Ave” and “Min” binding specificity metric due to mutation before and after some critical KiH (ie ScP, steric-complementary pair) swaps are applied to IgG1 Fc domain. A specificity metric change was derived from changes in affinity calculated using the “PathAve_DNN” stability change metric applied to all protein species, (complex, ligand, receptor). Here 200 mutation paths were averaged over for each calculation of the stability change metric.

To compare with “physical” 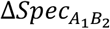, we passed the HetFc designs/mutations through our structural repacking workflow (Section S.2). Since the lab samples, corresponding to these mutations, were at least 85 % pure in the Fc Heterodimer, we expected that the “Ave” and “Min” 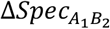 would be negative in value for most mutations in the HetFc dataset. However, like their DNN-based counterparts, only the physical “Ave” 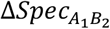, and not the physical “Min” 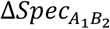, appeared to meet this expectation. Despite this, the distribution of “Min” 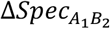 (Figure S7) are roughly centered about a value of zero, so there are favourable, negative-valued metric changes. Figure S8 shows that the PathAve_DNN, Local_DNN, and Global_DNN “Ave” 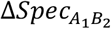 correlate well to the physics-based Amber and Electrostatic (Electro) “Ave” 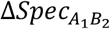. The PathAve_DNN and DDRW “Min” 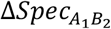 metrics correlate reasonably well with a Pearson r coefficient of ∼0.35; in addition, the negative predictive value (NPV) for predictions by the PathAve_DNN “Min” 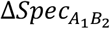 (of a favourable/negative-valued change) given the DDRW-based “Min” 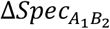 as truth is ∼0.76, meaning that 76 percent of the mutations deemed favourable by the PathAve_DNN “Min” 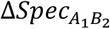 will also be deemed favourable by the DDRW-based “Min” 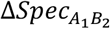.

### 5.4 Automated HetFc Design on a collection of target regions

We ran automated design optimization on each of the 74 regions associated with mutations in our HetFc stability dataset. The starting, test structure was again the Fc domain (PDB id, 3AVE) of an IgG1 antibody. For each region, the 150 sequence snapshots generated were hierarchically clustered. For a given target region, clusters having 4 or more sequence snapshots were considered; in particular, we focussed on the 2 clusters which had the lowest average value of the Local_DNN specificity metric, 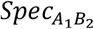. Unique mutations (ie, sequences) from each of these 2 clusters for every target region were collected and pooled together, then all repacked via our structural repacking workflow (Section 4.2). We then compared the change in the “Ave” and “Min” specificity metrics due to mutation with respect to wild-type, 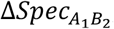, for all generated mutations/designs for several different choices of underlying stability metric: standard “physical” metrics (Amber, Electro, LJ, and DDRW) as well as DNN-based metrics (Local_DNN, Global_DNN, PathAve_DNN, and Decouple_DNN). Simulation trajectories were generated using two protocols: one with W = 1.0, η = 0.25 (“wt_1.0_temp_0.25”, strong bias); alternatively, one with W = 0.0, η = 1.0 (“wt_0.0_temp_1.0”, no bias). When W = 0.0, the Metropolis condition is turned off and so swaps drawn from the conditional probability distribution (ie, ZymeSwapNet output) are immediately accepted. The procedure above yielded 1302 and 2273 unique mutations (ie, Fc heterodimer *A*_1_*B*_2_ target region sequences) for the “wt_0.0_temp_1.0” and “wt_1.0_temp_0.25” sets of simulation runs, respectively. Swaps to CYS and PRO were not allowed.

After a switch in simulation protocol from “wt_0.0_temp_1.0” (bias off) to “wt_1.0_temp_0.25” (bias on), the 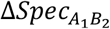 distributions from the generated designs deform consistently for the different choices of underlying stability metric (Figures 6, S9, S10, S11, and S12). Exact duplicates of a given *A*_1_*B*_2_ sequence/design were removed from the collection of sequences prior to determining a distribution. As hoped, the deformations of the “physical” 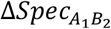 distributions matched the deformations of the DNN-based 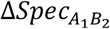 distributions: the distributions drifted to more favourable, negative values of the “Ave” (Figures S9 and S10) and “Min” (Figures 6, S11, and S12) 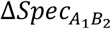. We expected that after the specificity bias was activated and the artificial temperature lowered, the distributions would deform, reflecting an increase in the number of HetFc selective designs. Although information from physical metrics was not used in the simulation method to generate the *A*_1_*B*_2_ sequences/designs, the distributions of the physical 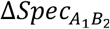 respond to the switch in protocol in the same manner as the distributions of the DNN-based 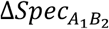. When the specificity bias was turned on, the correlations between DDRW-based “Min” 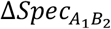 and the DNN-based “Min” 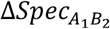 increased with the Pearson correlation coefficient going from ∼0.1 (bias off) to ∼0.47 (bias on); in addition, the negative predictive value (NPV) for predictions by the DNN-based “Min” 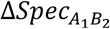 (of a favourable, ie, negative-value) given the DDRW-based “Min” 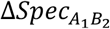 as truth increased from ∼0.1 to ∼0.4. With the bias on, when we re-defined the boundary between a favourable and an unfavourable 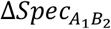 value to the 30^th^ percentile (an arbitrary choice), then this NPV measure of agreement between the DNN-based and the DDRW-based “Min” 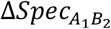 increased further to ∼0.7, meaning that 70 percent of those mutations deemed favourable by the DNN-based 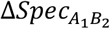 could also be deemed favourable by the DDRW-based 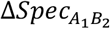.

**FIGURE 6.**
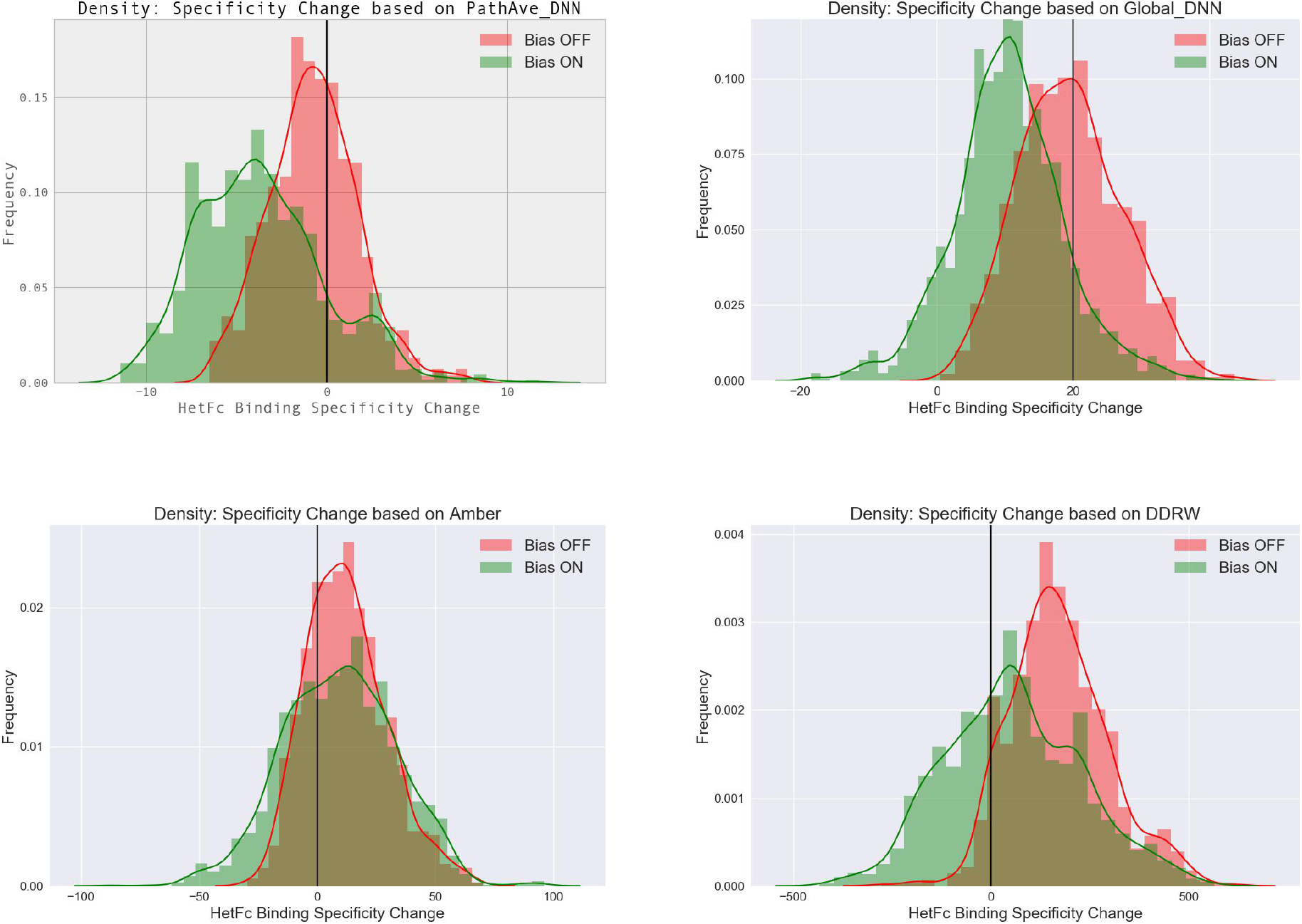
“Minimum” (Min) Specificity Metric change distributions from mutations generated by running automated sequence design on 74 target regions taken from our HetFc stability dataset. Binding specificity metric changes for two simulation protocols: wt_0.0_temp_1.0 (Bias OFF); wt_1.0_temp_0.25 (Bias ON). To evaluate the “PathAve_DNN” stability change metric for either a complex, ligand or receptor, 200 mutation paths were averaged over for each calculation of the stability change metric. Physics-based (Amber) and knowledge-based (DDRW) energy functions/stability metrics were calculated given structures output by our structural repacking workflow. Refer also to Figures S11 and S12.

### 5.5 Automated HetFc Design on two example Target Regions

We examine the *A*_1_*B*_2_ HetFc sequences generated by running automated design optimization on two example target regions (Table 3) in the HetFc. The first target region (“R1”) is taken from Ref 29 and corresponds to the mutation region associated with the top HetFc design engineered there using a KiH approach. The second region (“R2”) is a small expansion on R1. Both regions include the critical KiH positions taken from Ref 29: A/405 and A/407 (a “Hole” was created in Ref 29 with swaps A/405F → A and A/407Y → Y), as well as the corresponding position B/394 across the CH3 interface (a “Knob” was created in Ref 29 with swap B/394T → W). The exact KiH design from Ref 29 will not necessarily be generated from a single automated design run; however, given different choices in simulation protocol and simulation length, a diversity of different HetFc designs can be produced for any given target region. Four protocols were adopted: “wt_0.0_temp_1.0”, “wt_1.0_temp_1.0”, “wt_1.0_temp_0.5”, and “wt_1.0_temp_0.25”. The protocols in this list are ordered by increasing tendency to drive the HetFc system towards sequence states that favour the selective binding of the HetFc over the two corresponding Fc homodimers. Swaps to CYS and PRO were omitted from the sampling for target region “R1”, but allowed for “R2”. For any given simulation run, 150 sequence snapshots were collected and clustered hierarchically, generating dendrograms.

**TABLE 3.**
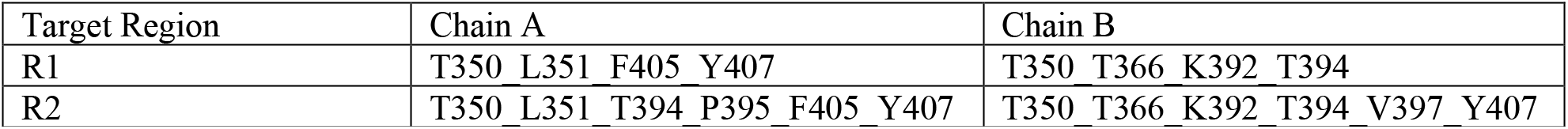
Example target regions on Fc domain (PDB id, 3AVE) for automated specificity optimization and HetFc design.

| Target Region | Chain A | Chain B |
| --- | --- | --- |
| R1 | T350 L351 F405 Y407 | T350 T366 K392 T394 |
| R2 | T350 L351 T394 P395 F405 Y407 | T350 T366 K392 T394 V397 Y407 |

In Figures 7 and S13 we show dendrograms derived from clustering R1 sequences generated by three separate simulation runs corresponding to three different simulation protocols. As the specificity bias increased, the number of distinct clusters decreased (Figure S13) and also the distributions of the “Ave” and “Min” 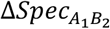 shifted to more favourable, negative values (Figures 7, 8, S14, and S15). Again although physical metrics were not used in the automated design to generate *A*_1_*B*_2_ sequences, extracting the unique sequences and running them through a structural repacking workflow yielded reasonable changes to the physical 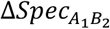 (Figures 8 and S15). Correlations became strong between the DNN-based and physical “Ave” 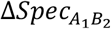 once the simulation protocol became sufficiently strong at driving a selective binding towards the HetFc (Figure S16). For the “wt_1.0_temp_0.5” protocol, we saw a recognizable Knobs-into-Holes (KiH) design with critical swaps, *A*_1_ swap A/407Y → G and *B*_2_ swap B/394T → Y (Tables 4 and 5), with a very favourable, negative value in the change of the DDRW-based “Min” 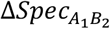 (Table 5). In addition, there were electrostatic steered designs with critical swaps, *A*_1_ swap A/407Y → E and *B*_2_ swap B/394T → H: here the favourable Fc heterodimer formation was driven by a local electrostatic attraction, and one of the Fc homodimers, *A*_2_*B*_2_, had a relatively unfavourable “Knob-Knob/clashing” interaction between wild-type residue A/407Y and the *B*_2_ swap B/394T → H. The other swaps at the remaining *A*_1_*B*_2_ target residue positions appeared very similar to those chosen by in Ref 29 to meet a goal of stabilizing the HetFc design : our automatically generated swaps were either identical or similar to the stabilizing *A*_1_ swaps (A/350T → V, A/351L → Y, A/405F → A) and *B*_2_ swaps (B/350T → V, B/366T → L, B/392K → L) lifted from Ref 29. Tables 4 and 5 list two of the most selective designs. Table 4 also gives the absolute binding affinities for all three Fc complexes (*A*_1_*B*_2_, *A*_1_*B*_1_, *A*_2_*B*_2_) calculated using the Global_DNN energy metrics. The unfavourable “Knob-Knob/clashing” interaction is clearly indicated by the large increase in the value of the Global_DNN binding affinity of the homodimer *A*_2_*B*_2_ going from the wild-type design to the HetFc selective design.

**FIGURE 7.**
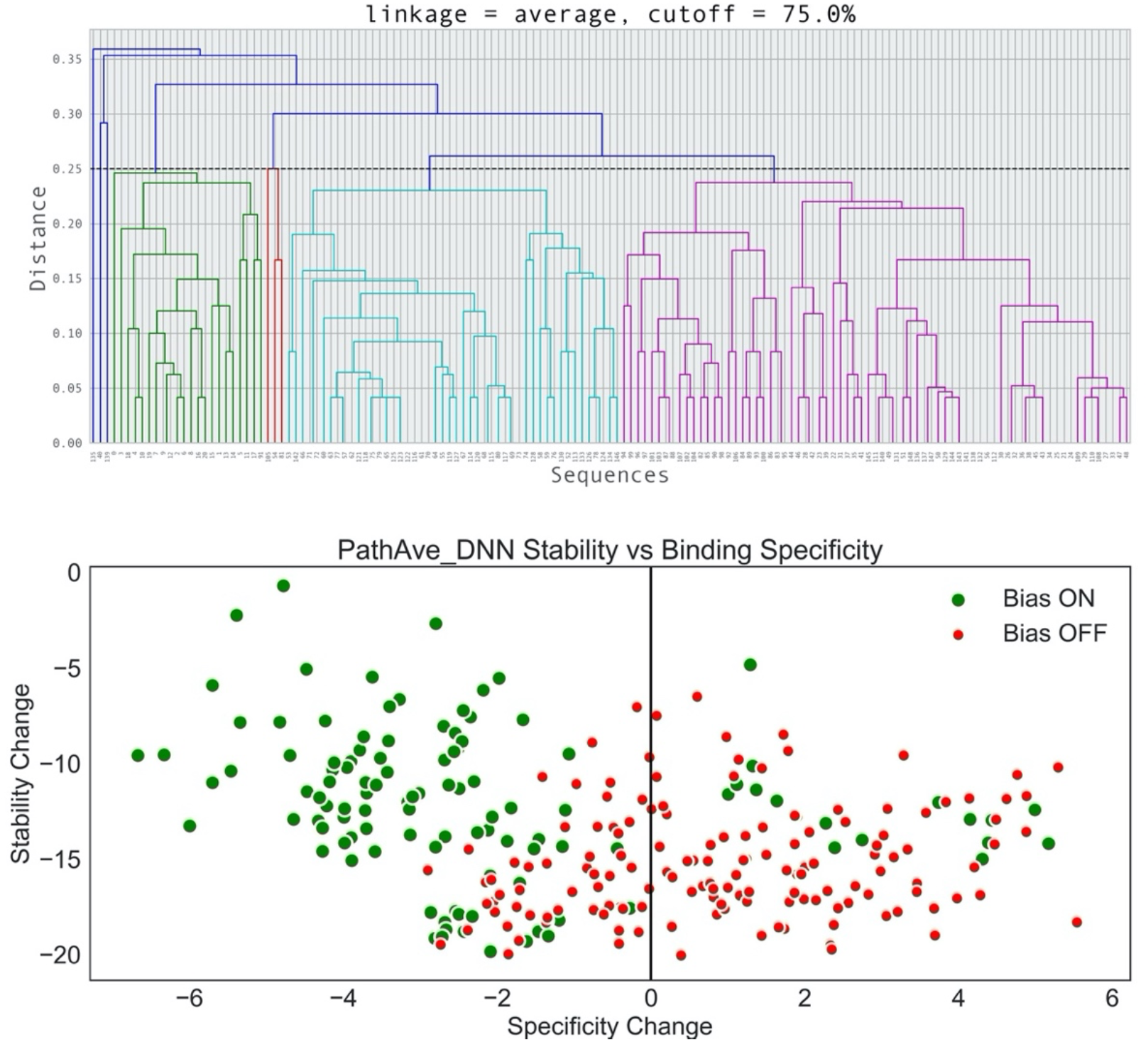
Simulation trajectory sequence snapshots generated by an automated specificity design run on the R1 target region of the Fc domain. For a given simulation run 150 simulation snapshots were generated, then a dendrogram was determined by hierarchical sequence clustering these snapshots. The sequence similarity cutoff was 75 percent: all sequences in a cluster (same colour *other than blue*) therefore were at least 75 percent similar. Top panel, dendrogram for simulation protocol “wt_1.0_temp_0.5” (Specificity Bias ON): cyan branch, electrostatically driven designs; purple branch, sterically driven (‘KiH’) designs. Bottom panel, change in stability metric vs change in “Min” specificity metric due to mutations generated from two simulation protocols, “wt_0.0_temp_1.0” (red points, Specificity Bias OFF) and “wt_1.0_temp_0.5” (green points).

**FIGURE 8.**
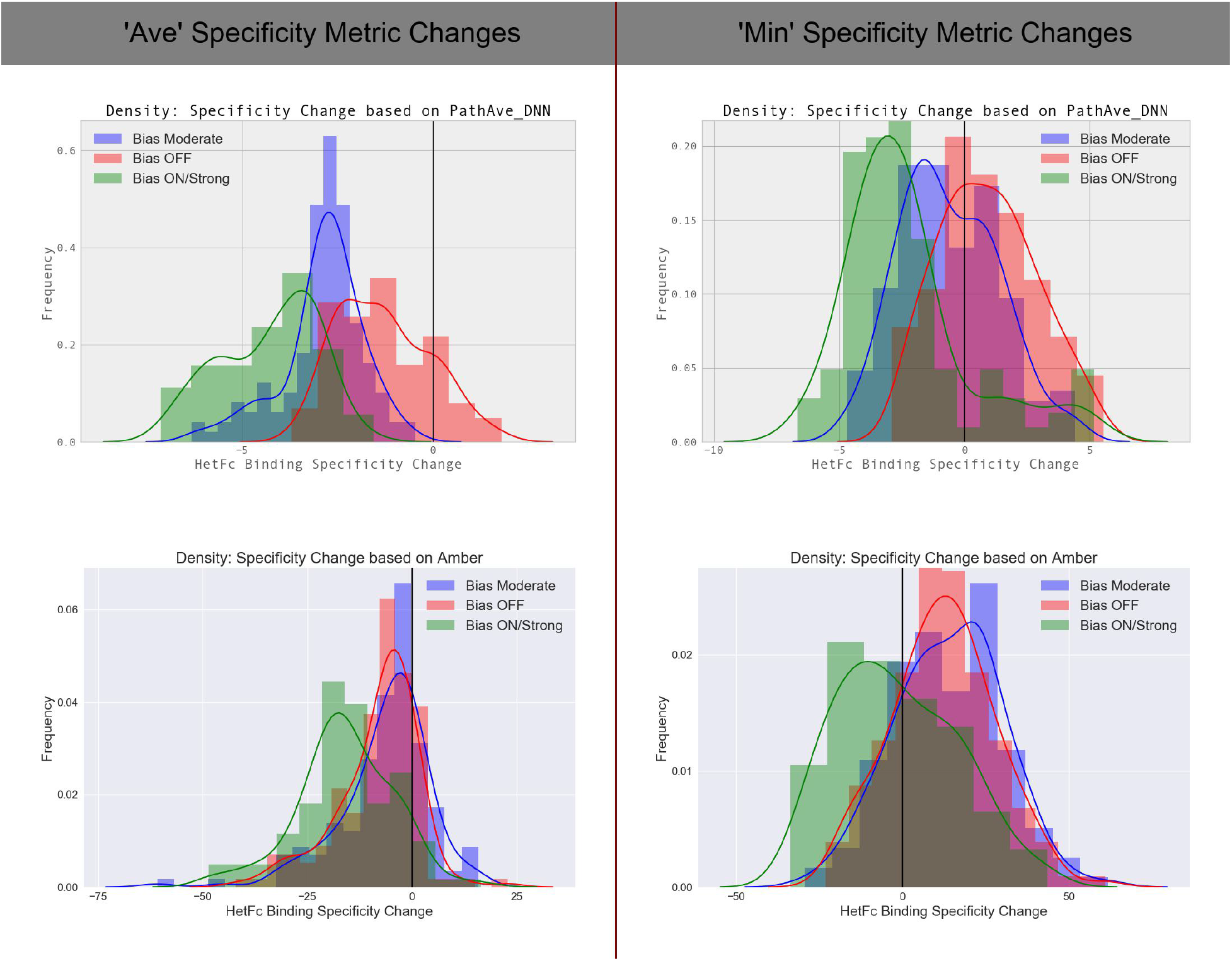
Distributions of the change in the specificity metrics due to mutation relative to wild-type for sequences generated by an automated specificity design run on the R1 target region in the IgG1 Fc domain.. Three simulation runs were performed each following a different protocol: (Red) wt_0.0_temp_1.0; (Blue) wt_1.0_temp_1.0; and (Green) wt_1.0_temp_0.5. Only unique *A*_1_*B*_2_ sequences / designs were used when determining a distribution: there were no repeated sequences. To evaluate the “PathAve_DNN” stability change metric for either a complex, ligand, or receptor, 200 mutation paths were averaged over for each calculation of the stability change metric. Specificity metric changes based on the physics-based (Amber) energy function/stability metric were calculated on structures output by our structural repacking workflow. Refer also to Figures S14 and S15.

**TABLE 4.** Absolute binding affinities for the Fc heterodimer and homodimers. Shown are four of the most HetFc selective binding sequences generated by an automated specificity design run on the R1 target region. Swaps to CYS and PRO were not allowed in this run. The wild-type reference design is colored in black and highlighted in bold. Note the following sign convention: as a binding affinity becomes more negative in value, the binding itself becomes more favourable. Simulation protocol was “wt_1.0_temp_0.5” (see discussion in Section 5.5). Second row, electrostatically driven design; third row, sterically driven (‘KiH’) design.

| HetFc Design $A_1B_2$ , Target Region R1 | Global_DNN<br>$A_1B_2$ | Global_DNN<br>$A_1B_1$ | Global_DNN<br>$A_2B_2$ |
| --- | --- | --- | --- |
| A/350T_A/351L_A/405F_A/407Y_B/350T_B/366T_B/392K_B/394T | -14.644 | -14.644 | -14.644 |
| A/350V_A/351F_A/405S_A/407E_B/350A_B/366G_B/392I_B/394H | -12.538 | -10.865 | -2.122 |
| A/350V_A/351F_A/405S_A/407G_B/350I_B/366G_B/392L_B/394Y | -19.178 | -18.817 | -2.434 |

**TABLE 5.** “Min” Binding Specificity Metric change for the Fc heterodimers. A Knobs-into-Holes (KiH) and a disulphide HetFc selective binding sequences generated by automated specificity design run on the R1 target region. Swaps to CYS and PRO were not allowed. Simulation protocol was “wt_1.0_temp_0.5” (see discussion in Section 5.5). First row, electrostatically driven design; second row, sterically driven (‘KiH’) design.

| HetFc Design $A_1B_2$ , Target Region R1 | PathAve_DNN | Local_DNN | Global_DNN | Amber | DDRW |
| --- | --- | --- | --- | --- | --- |
| A/350V_A/351F_A/405S_A/407E_B/350A_B/366G_B/392I_B/394H | -3.624 | -4.702 | -1.673 | -26.36 | -37.43 |
| A/350V_A/351F_A/405S_A/407G_B/350I_B/366G_B/392L_B/394Y | -6.671 | -4.198 | -0.361 | 17.27 | -219.43 |

Allowing for swaps to CYS and PRO, the automated specificity design on region R2 generates a HetFc selective design very similar to the KiH design of Ref 29 with a TRP at sequence position B/394 (*B*_2_ swap B/394T → W) and a GLY at A/407 (*A*_1_ swap A/407Y → G). In addition, for this design, the HetFc binding is further strengthened by the charge-polar interaction between the GLU at A/395 and a SER or ARG at B/397 (Tables S4 and S5). This KiH design was generated using the “wt_1.0_temp_0.25” simulation protocol. Along the simulation trajectory the TRP at B/394, the GLY at A/407, and a LEU at B/407 were conserved, while the remaining target region positions showed some fluctuations in amino acid identity. To close, we emphasize that the automated specificity design method is stochastic; consequently, different runs under the same simulation protocol can produce a different ensemble of *A*_1_*B*_2_ sequence designs, but collectively these ensembles have the advantage of providing a diversity of optimal HetFc sequence solutions.

## 6 Summary

ZymeSwapNet was trained to predict the amino acid identity at an arbitrary target residue site in a protein given a simple coarse-grained protein representation as input features, which includes the amino-acid identities of residues neighbouring the target but excludes the side-chain atomic conformations/coordinates. Compared to other state of the art neural network models, ZymeSwapNet’s accuracy/performance is comparable, despite its simplicity. Compared to traditional computational protein design methods, which utilize force-fields, ZymeSwapNet’s accuracy in wild-type recovery tests is about 10 to 15 percent higher. Although ZymeSwapNet was trained to effectively predict a protein sequence (ie, by applying ZymeSwapNet repeatedly across residue sites), it can also be used to target various design goals such as to discover potentially stabilizing swaps. For the IgG1 Fc domain structure, which was re-engineered into a Fc heterodimer [25, 29], ZymeSwapNet correctly recommends some of the same stabilizing swaps found necessary to build-back stability lost during the earlier negative design rounds [29]. In the first stage of that effort, Fc stability was lost after the applying the Knobs-into-Holes (KiH) swaps, which were critical for HetFc specific pairing. The stabilizing swaps, complementing the KiH swaps, were predicted by ZymeSwapNet to be more probable than the corresponding wild-type amino-acid identities. This finding suggested that the probability distributions output by ZymeSwapNet could be used to formulate stability and affinity metrics [38-40]. In this paper, ZymeSwapNet derived affinity metrics were tested on a subset of mutations listed in the open-source protein-protein affinity database, SKEMPI [41]. Affinity predictions based on ZymeSwapNet metrics correlated well with experimental protein binding affinity measurements, such that the degree of this agreement was comparable to that exhibited by other standard physics-based (Amber for instance) and knowledge-based (DDRW) affinity metrics. Unlike these physical metrics, the calculation of ZymeSwapNet metrics is fast because it does not require side-chain atomic coordinate information, that is it avoids lengthy side-chain conformational repacking; furthermore, as a result, it avoids being beleaguered by issues associated with repacking like the choice of force-field, the treatment of protonation state, and the potential deficiencies arising from the reliance on a rotamer library.

ZymeSwapNet can quickly rank and screen mutations, but it can also be used within an automated sequence optimization algorithm which rapidly proposes novel mutations/designs, potentially satisfying multiple protein design objectives. The only inputs to the sampling are a starting structure and the list of target residue sites which, although not examined in this paper, could in principle include all sites in the protein structure. There is no restriction to the selection of these residues: they do not have to be contiguous in protein sequence position, and the structure may contain multiple chains. An example is the design of a HetFc where the two objectives are the enhancement of HetFc stability and binding specificity. This stochastic Monte Carlo sampling algorithm, which incorporates the conditional probability represented by ZymeSwapNet to make instantaneous amino-acid assignments and a Metropolis-based binding specificity bias, was found capable of driving an ensemble of HetFc designs towards stronger binding specificity, and for some target region selections rapidly generating Knobs-into-Holes HetFc designs very similar to those discovered through multiple rounds of rational design and experimental verification [29] as well as novel HetFc designs. We have demonstrated that such deep learning-based algorithms will prove very valuable in addressing multi-objective protein design optimization problems, generating a diverse yet focussed ensemble of potential designs.

## Acknowledgments

The authors would like to thank Nicholas Geraedts, Charles M. Stevens, and Dimitri Tcaciuc for their helpful discussions and technical support on this work.

