## Supplementary Information for "Automated Protein Affinity Optimization using a 1D-CNN Deep Learning Model"

**S.1 Model Training**

**S.2 Scoring Mutations using ZymeSwapNet**

**S.3 Automated Binding Specificity Optimization**

**S.3.1 Automated HetFc Design on a collection of target regions**

**S.3.2 Automated HetFc Design on two example Target Regions, R1 and R2**

**S.4 References**

### S.1 Model Training

Based on PDB id, we divided a culled PDB features dataset first into a development set (80 % of the PDB ids) and a hold-out set (20 % of the PDB ids), then "fold" the development set into 5 possible (training set, validation set) combinations to cross-validate model training. In Table S1, we report the average accuracies at predicting the standard one-letter-code amino-acid identities (ie, 20 classes) at target residue sites calculated over the validation sets of a 5-fold partitioning of the development set taken from the pc90 culled PDB features dataset. In addition, we report the prediction accuracies on the hold-out set where ZymeSwapNet was retrained on data from the entire development set.

| Features | Fold-Type | Top-1 Acc. | Top-2 Acc. | Top-3 Acc. | Top-4 Acc. | Average Sensitivity | Average Precision |
|---|---|---|---|---|---|---|---|
| BB | Fold(s) | 0.468 (0.001) | 0.627(0.001) | 0.716(0.001) | 0.776(0.001) | 0.404 | 0.453 |
| BB | Hold-out | 0.477 | 0.636 | 0.724 | 0.784 | 0.414 | 0.469 |
| BB + SC | Fold(s) | 0.524(0.002) | 0.684(0.002) | 0.766(0.001) | 0.819(0.001) | 0.461 | 0.501 |
| BB + SC | Hold-out | 0.533 | 0.693 | 0.774 | 0.826 | 0.472 | 0.515 |

**TABLE S1: Cross-validation Top-N classification accuracies and Top-1 Sensitivity and Precision.** Quantity in brackets is the standard deviation evaluated over 5-folds of the (AU, pc90) development set. Number of data points in the development dataset was roughly 1,800,000, while 350,000 points were in the hold-out dataset; within each fold of the development dataset, roughly 1,500,000 points were in a training set while 350,000 points were in a validation set. With BB + SC features, a data point/object had a shape ((K = 19), 40) ; with BB features, a data point/object had a shape ((K = 20), 38). Here K denotes the number of neighbouring residues to a target residue site.

To quantify how well the trained neural network predicts different amino-acid identities, we calculated the precision and sensitivity metrics, along with the F1-score, associated with each amino acid identity. Here for the calculation of a metric for a given amino-acid identity ($RES$), we considered a binary classification problem:

- if the amino-acid identity, $RES$, is predicted (ie, the output node with the highest probability corresponds to $RES$) and the target residue label is also $RES$, then we have a true positive and so accumulate the $TP$ count;
- if the network predicts $RES$ but does not match the target residue label, then we have a false positive and so accumulate the $FP$ count;
- if the network predicts a amino-acid identity other than $RES$ but the target residue label is RES, then we have a false negative and so accumulate the $FN$ count.

Precision and sensitivity are defined in the usual way, $Prec = TP/(TP + FP)$ and $Sens = TP/(TP + FN)$. Sensitivity is the percentage of wild-type target residue labels, ie, amino-acid identities, that are are predicted correctly, while precision is the percentage of predictions that are correct. The F1-score is the harmonic mean of the sensitivity and the precision (Figure S2).

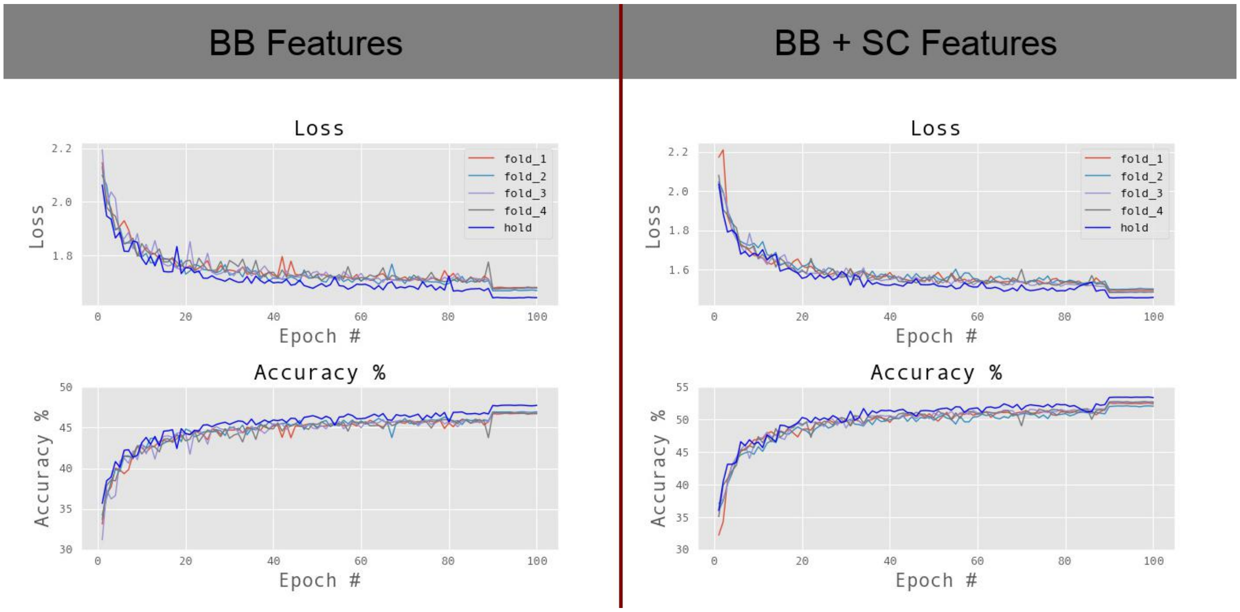

**FIGURE S1 Loss and accuracy versus epoch number for validation and hold-out sets.** Features dataset (AU, pc90)**.** 'fold_1' to 'fold_4' refer to 4 of the 5 cross-validation folds of the development set while 'hold' refers to the training on the entire development set, testing the amino-acid predictions on the hold-out set. Note adding a dropout of 1.5 % to the 1D-CNN sub-network and switching from a 'reLU' to an 'eLU' activation in this sub-network (Figure 1) increased the hold-out accuracy to 49% for BB features and 55% for BB+SC features (not shown).

An iterative, deterministic algorithm for predicting wild-type sequence, which utilizes the two ZymeSwapNet models, BB and BB + SC, should yield a wild-type sequence recovery accuracy bounded between roughly 47.7 % and 53.3 %. Starting with a protein backbone stripped of its sequence identity, such an algorithm could use the BB ZymeSwapNet model for the initial sequence assignment, followed by iterative applications of the BB + SC ZymeSwapNet model until sequence convergence. An application here would involve assigning the most probable amino-acid identity to the current target residue site visited, then communicating this assignment to residues having this target residue as one of their neighbours.

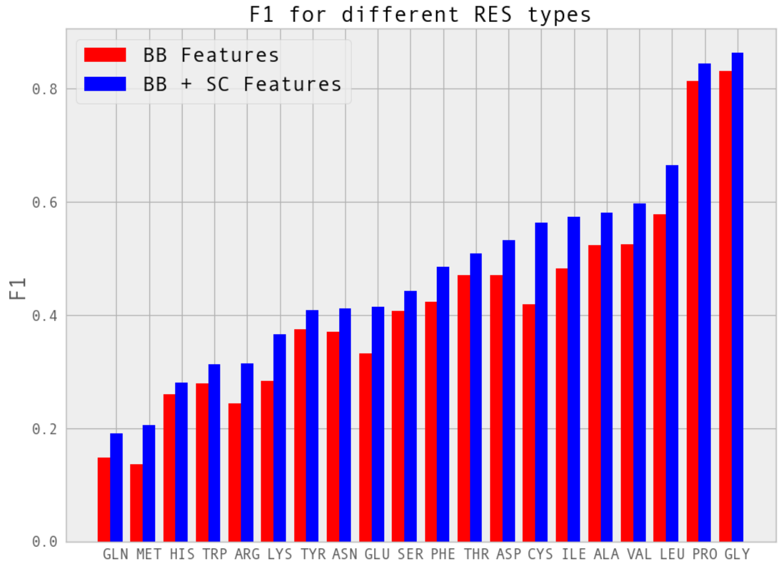

**FIGURE S2 F1-Score on hold-out set.** ZymeSwapNet DNN models were trained on entire development set, then the F1-Score was evaluated on the hold-out set.

PRO and GLY were predicted correctly with very high accuracy, because of their quite distinct Ramachandran Angle distributions and because GLY, being such a small residue, is characterized by a relatively tight cluster surrounding it - note one set of pair-residue features are the distances from the target residue site to the nearest-neighbouring residue sites. Hydrophobic residues, located typically in more buried, sterically hindered regions of PDB crystal structures, were predicted correctly with higher accuracy than the more charged residue types, which lie in the more sequence-variable surface-exposed regions.

Figure S3 shows the frequencies of predicted amino-acid identities as heat-maps. In the first plot, number 1, given the PDB amino-acid identity label at the target residue, we plot the frequencies of occurrence of the most-probable (ie Top-1) amino-acid identity predicted. In the second plot, number 2, given the identity of the most-probable amino-acid identity predicted, we plot the frequencies of occurrence of the second-most (ie, next-most) probable amino-acid identity predicted. Explicitly, these two frequencies for amino-acid identities $l$ and $p$ are

$$H_{l,p}^{(1)} = \frac{\sum_t I[l = RES_t] I\left[p = \mathrm{argmax_q}\left(\mathrm{Prob}(\mathrm{q}; t)\right)\right]}{\sum_{t'} I[l = RES_{t'}]}$$

$$H_{l,p}^{(2)} = \frac{\sum_t I\left[l = \mathrm{argmax_q}\left(Prob(\mathrm{q}; t)\right)\right] I\left[p = \mathrm{argmax_q}(\mathrm{Prob}(\mathrm{q}; t); 2)\right]}{\sum_{t'} I\left[l = \mathrm{argmax_q}\left(\mathrm{Prob}(\mathrm{q}; t')\right)\right]}$$

where $I[\cdot]$ is an indicator function, which equals 1 if the equality in its argument is satisfied, and equals 0 otherwise. The neural network at any given target residue site $t$ generates an output, discrete probability function with 20 values where each value corresponds to the probability $\mathrm{Prob}(q; t)$ of amino-acid identity $q$ at site $t$. The amino-acid identity at the site $t$, that labels that site, is $RES_t$. The $\mathrm{argmax_x}(\mathrm{Prob}(x; t); 2)$ returns the identity of the second-most probable amino-acid identity.

For Figure S3, the rows and columns of the $\boldsymbol{H}$ matrices were reordered so as to minimize the *bandwidth* of these matrices: here a sparse matrix $\boldsymbol{H}_{sparse}$ was generated by dropping elements of $\boldsymbol{H}$ whose magnitudes were less than some arbitrarily chosen threshold value; permutation operations that minimize the bandwidth of the symmetrized version of $\boldsymbol{H}_{sparse}$ were discovered via the *Cuthill-McKee algorithm* and then applied to the original matrix $\boldsymbol{H}$. Minimizing the bandwidth has the effect of clustering amino-acid identities along the axes of these two plots.

The heat-map plots of $H_{l,p}^{(1)}$ and $H_{l,p}^{(2)}$ reveal several groupings of predicted amino-acid identities. If one amino-acid identity is predicted from the group, then the other members of the group are quite likely to be predicted at the target residue site $t$ :

1. (TYR, PHE);
2. TRP→ (TYR, PHE);
3. (ILE, VAL);
4. (ASN, ASP);
5. (GLN → GLU);
6. (LYS, ARG); and
7. (SER, THR).

When we write A→ B here, we mean the following: if A is predicted then B is predicted as also a likely amino-acid identity, but if B is predicted then A is not predicted as a likely amino-acid identity.

<table>
<tr><th>Features</th><th>1, $H_{l,p}^{(1)}$</th><th>2, $H_{l,p}^{(2)}$</th></tr>
<tr><td>BB</td><td>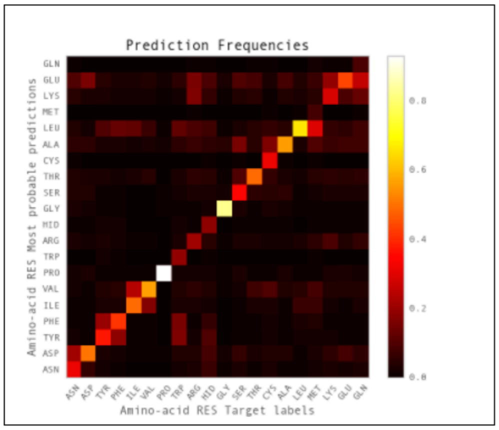
</td><td>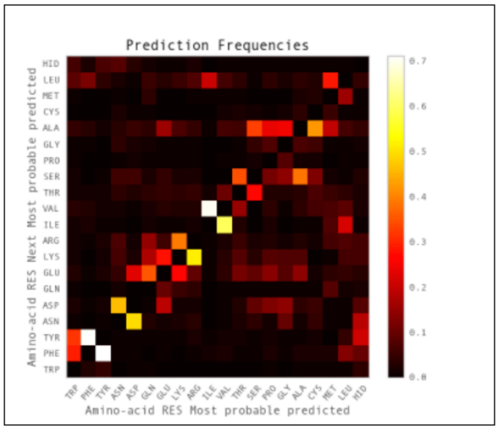
</td></tr>
<tr><td>BB + SC</td><td>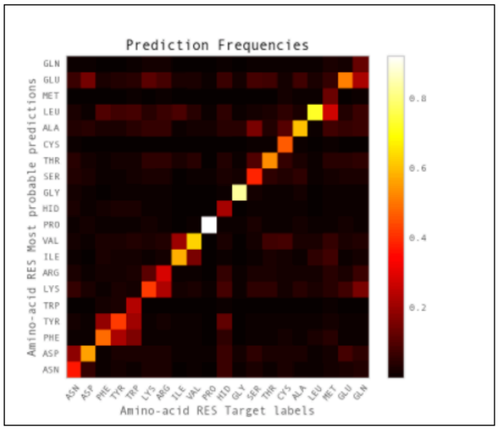
</td><td>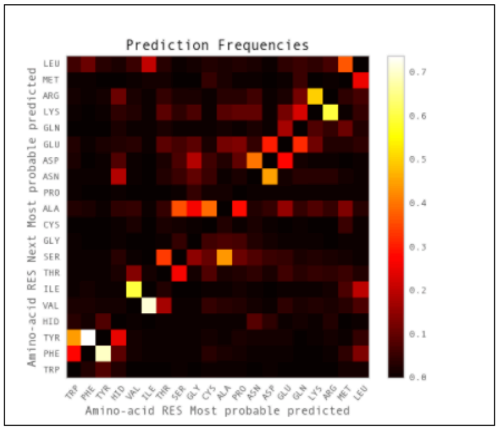
</td></tr>
</table>

**FIGURE S3 Predicted Amino-acid Frequencies of occurrence**

#### S.2 Scoring Mutations using Deep Neural Networks

##### Structural Repacking Workflow

We score protein designs/mutations. DNN based scores do not require information on side-chain conformation; however, we do make comparisons to traditional, "physical scoring metrics" (Amber, LJ, Electro, and DDRW) and to evaluate these we do require this information. To evaluate changes to "physical" metrics due to mutation, the side-chains in a local region surrounding the swap sites are repacked to accommodate the new amino-acid identities introduced into the structure. In order to minimize the Amber energy, our repacking method performs local conformational changes, optimizing the protein conformation in a local, "mobile" region, surrounding and including the mutation/swap sites. Our repack has two stages. The first stage is a cyclic steepest descent in rotamer space given a fixed protein backbone [1], otherwise known as "Iterated conditional modes" (ICM) [2]. Our standard side-chain rotamer library is the "honig-5" library, a very large amino-acid identity specific library prepared by Honig group at Columbia University [1]. In the first stage, 20 independent steepest descent runs are performed: each run starts from a common initial condition where all residues in the mobile region, with the exception of those at the swap sites, have their wild-type crystal conformations, while an arbitrary rotamer from the library is assigned to the residues at the swap sites. Each of the 20 runs then optimizes the residues in the mobile region in a random order, making each run distinct from any other. Once all 20 runs are complete, the lowest energy structure is output and fed into the final stage of the repack. In the final stage the constraint of a fixed backbone is released. The sidechains and backbone of the mobile region are simultaneously co-optimized using a pair of deterministic minimizers: a quick steepest descent to remove atomic clashes, followed by L-BFGS minimization.

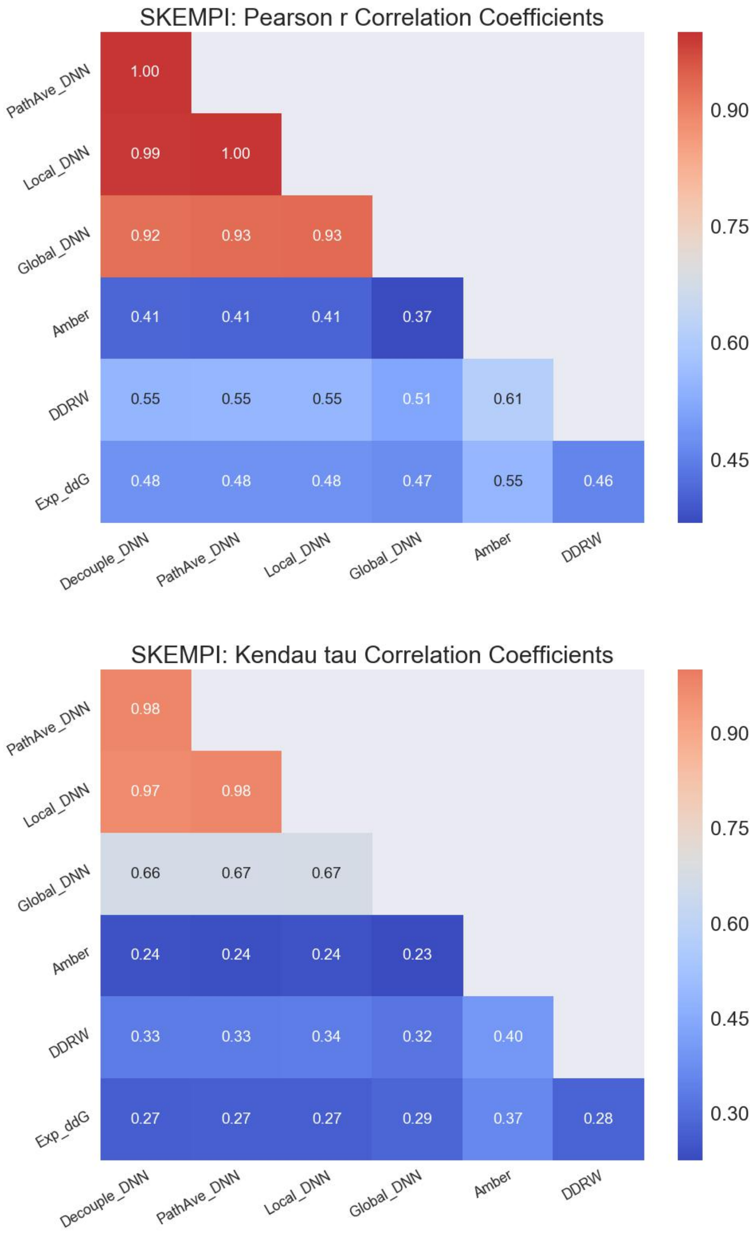

**FIGURE S4 Correlations (Pearson r and Kendall tau) between predictions (DNN-based, physics-based and knowledge-based metrics) and experimental measurements (SKEMPI dataset) of protein-protein binding affinity changes due to mutations from wild-type**. All DNN-based metrics employed neighbouring amino-acid identities) as features (ie, BB + SC, see Section 3). For the calculation of the "Decouple_DNN", swaps within any given mutation were treated as being decoupled: here when calculating the swap contribution to the stability change, residues neighbouring the swap site had their amino-acid identities set to wild-type. Correlation coefficients dropped significantly when a ZymeSwapNet model trained only on BB features was used to calculate DNN-based metrics.

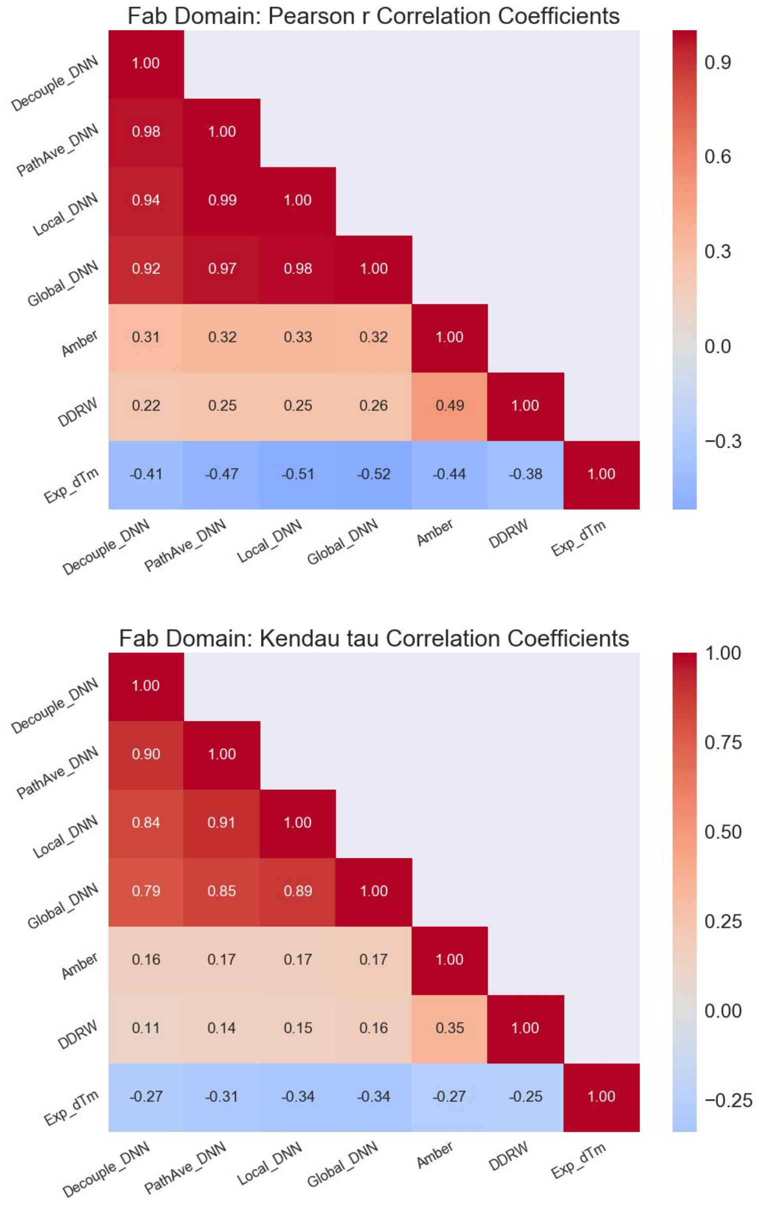

**FIGURE S5 Correlations (Pearson r and Kendall tau) between predictions (DNN-based, physics-based, and knowledge-based metrics) and experimental measurements (Exp_dTm, internal FAB dataset) of protein-protein stability changes due to mutations from wild-type.** All DNN-based metrics were evaluated from the 1D-CNN ZymeSwapNet model taking both backbone (BB) and side-chain (SC, neighbouring amino-acid identities) as features (ie, BB + SC, see discussions in Section 3). For the calculation of the "Decouple_DNN", swaps in any given mutation were treated as being decoupled: here when calculating the swap contribution to the stability change, residues neighbouring the swap site had their amino-acid identities set to wild-type. As expected all changes in the stability metrics due to mutation are anti-correlationed with Exp_dTm.

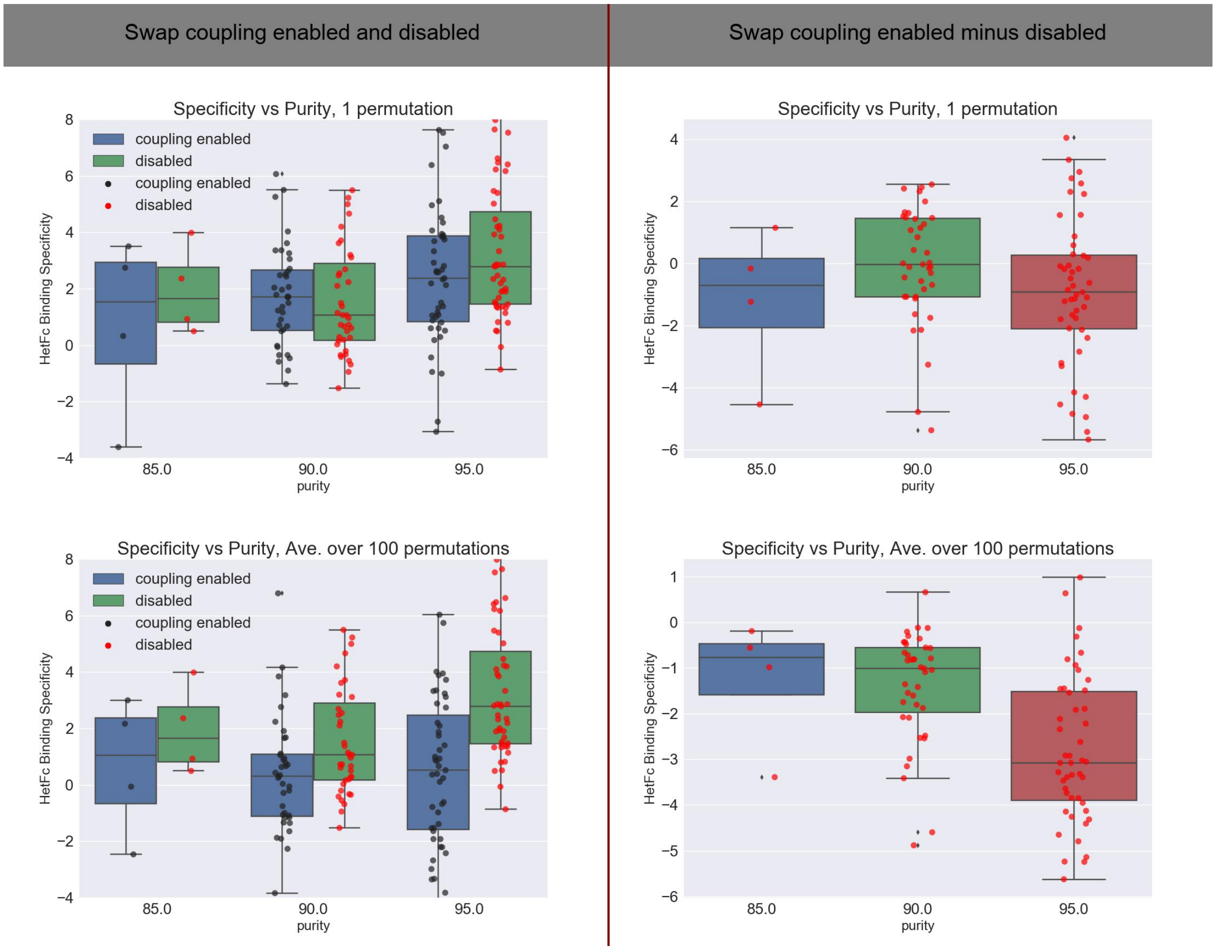

**FIGURE S6 "Min" Specificity metric changes due to mutation for mutations taken from our HetFc stability dataset.** Comparison of binding specificity changes between runs with and without coupling ("non-additive" effects enabled and disabled respectively) between multiple swaps in a mutation. "PathAve_DNN" stability change metrics were calculated from an average over a single (first row) or 100 (second row) mutation paths. The "Delta calculation" (second column) for a mutation is the binding specificity metric evaluated with coupling between swaps minus the binding specificity metric evaluated without coupling between swaps.

| Dimer Type | KiH | Complex | Mutation String |
|---|---|---|---|
| Hetero | On | $A_1B_2$ | A/397.VAL->SER A/405.PHE->ALA A/407.TYR->VAL B/392.LYS->VAL B/394.THR->TRP |
| Homo | On | $A_1B_1$ | A/397.VAL->SER A/405.PHE->ALA A/407.TYR->VAL B/397.VAL->SER B/405.PHE->ALA B/407.TYR->VAL |
| Homo | On | $A_2B_2$ | A/392.LYS->VAL A/394.THR->TRP B/392.LYS->VAL B/394.THR->TRP |
| Hetero | Off | $A_1B_2$ | A/397.VAL->SER A/407.TYR->VAL B/392.LYS->VAL |
| Homo | Off | $A_1B_1$ | A/397.VAL->SER A/407.TYR->VAL B/397.VAL->SER B/407.TYR->VAL |
| Homo | Off | $A_2B_2$ | A/392.LYS->VAL B/392.LYS->VAL |

**TABLE S2 Example of the removal of a KiH set of swaps from a HetFc mutation/design.**

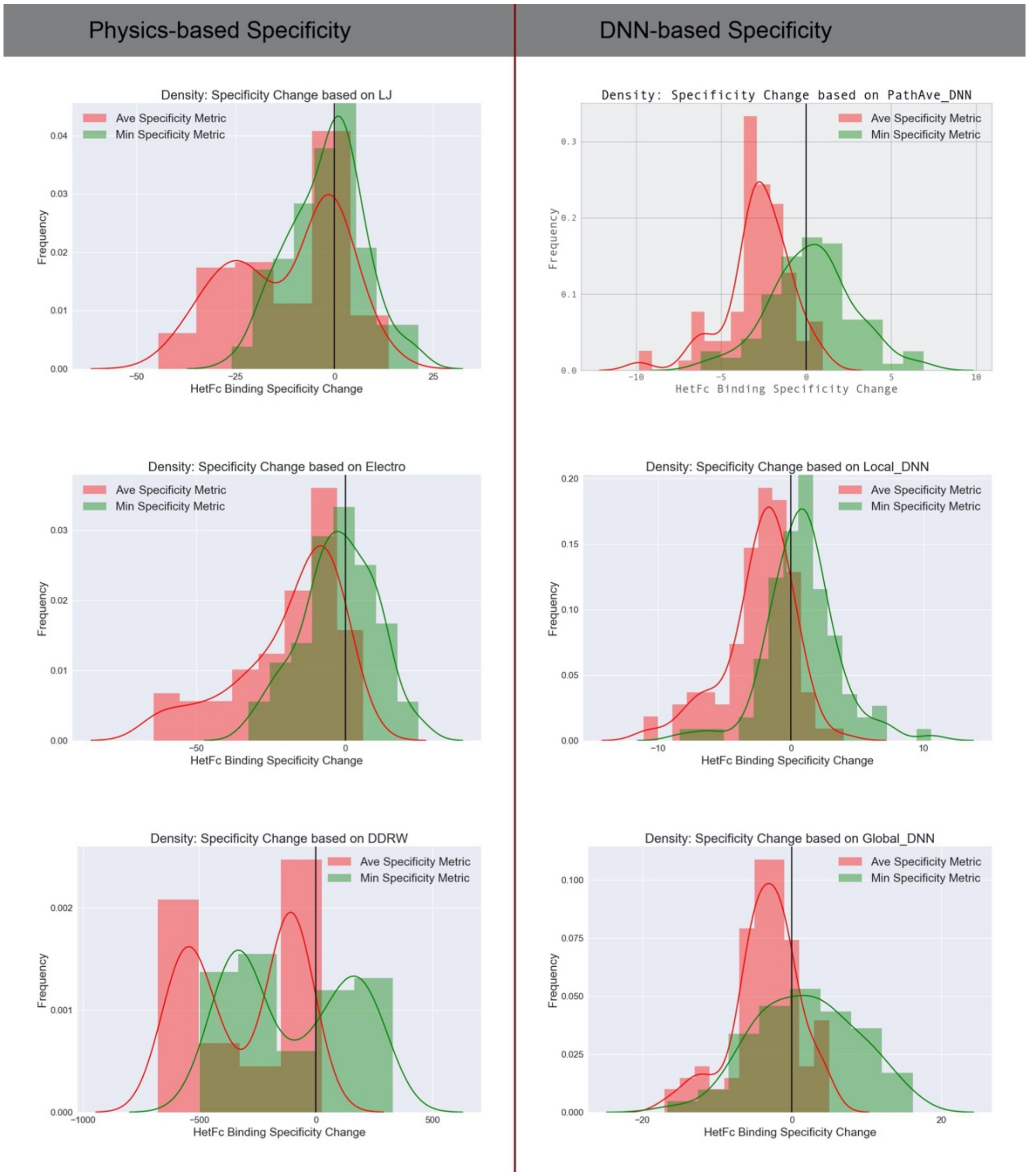

**FIGURE S7 "Average" (Ave) and "Minimum" (Min) Specificity Metric changes due to mutation for mutations taken from our HetFc stability dataset.** Binding Specificity Metrics were evaluated from a selection of physics-based and DNN-based energy functions. To evaluate the "PathAve_DNN" stability change metric for either a complex, ligand or receptor, 200 permutations / mutation paths were averaged over for each calculation of the stability change metric.

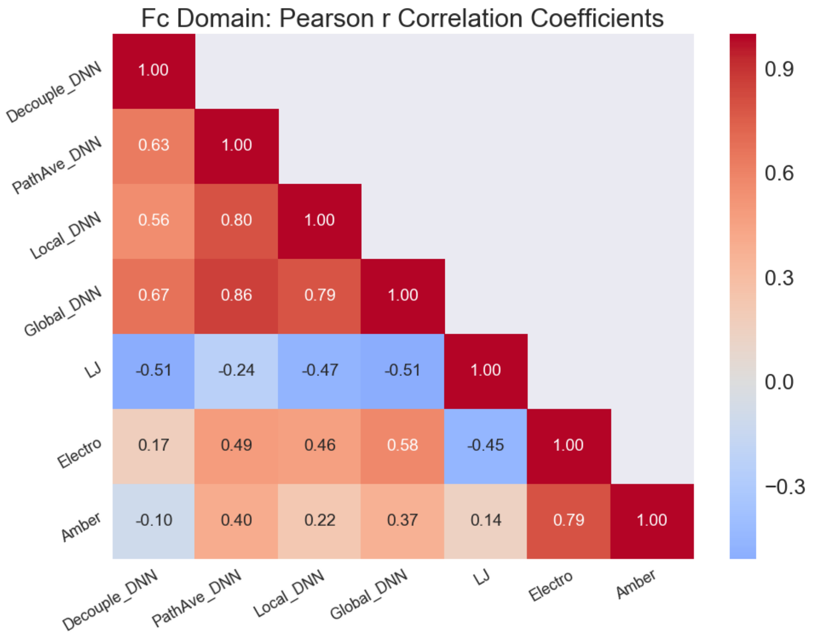

**FIGURE S8 Pearson r correlation coefficients between different "Average" (Ave) Binding Specificity Metrics changes due to mutation for mutations taken from our HetFc stability dataset.**

#### S.3 Automated Binding Specificity Optimization

Sequence similarity scores, required in order to assign cluster identities for a specified target region, were calculated based on a "fine-grained" grouping of amino-acid identities. For two $A_1B_2$ Fc heterodimer sequences being compared, if at a given target site the two amino-acid identities exactly match, then the count, $C$ , increases by $\Delta C = 3$; if the two amino-acid identities are similar, meaning that they are in the same "fine-grained" amino-acid group (Table S3), then $C$ increases by $\Delta C = 2$; otherwise, $C$ increases by $\Delta C = 1$. The distance between two sequences is then 1 - $\sum_{i=1}^{T} \Delta C\left(seq_{t_i}^1, seq_{t_i}^2\right)/(3T)$ where $T$ is the number of positions in the sequence/mutation, ie, the number of residues in the target region, and $seq_{t_i}^1$ is the amino-acid identity at position/target site $t_i$ for sequence '1'. Two simulation snapshots could have the same $A_1B_2$ HetFc sequence, meaning a sequence similarity/distance equal to one/zero.

| Group | Amino-Acid Types |
|---|---|
| **Aliphatic** | A, V, I, L, M |
| **Aromatic** | F, Y, W |
| **Negative** | E, D |
| **Positive** | K, R, H |
| **Neutral Polar** | S, T, Q, N |
| **Other** | G, P, C |

**TABLE S3 Fine-grained grouping of amino-acid identities for Heterodimer sequence clustering.**

##### S.3.1 Automated HetFc Design on a collection of target regions

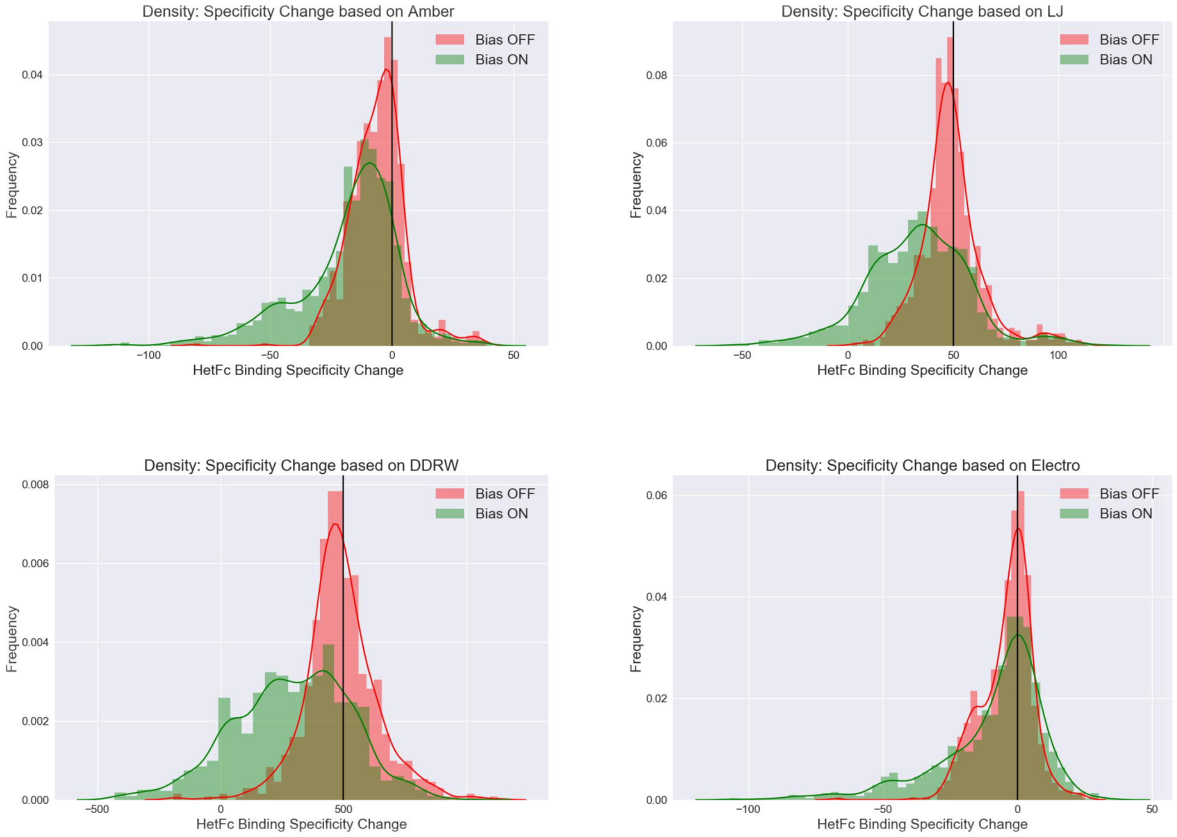

**FIGURE S9 "Average" (Ave) Specificity Metric change distributions from mutations generated by running automated sequence design on 74 target regions taken from our HetFc stability dataset.** Binding specificity metric changes due to mutation with respect to wild-type were calculated using physics-based and knowledge-based energy functions/stability metrics on structures output by our structural repacking computational workflow. Simulation protocols: wt_0.0_temp_1.0 (Bias ON); wt_1.0_temp_0.25 (Bias OFF).

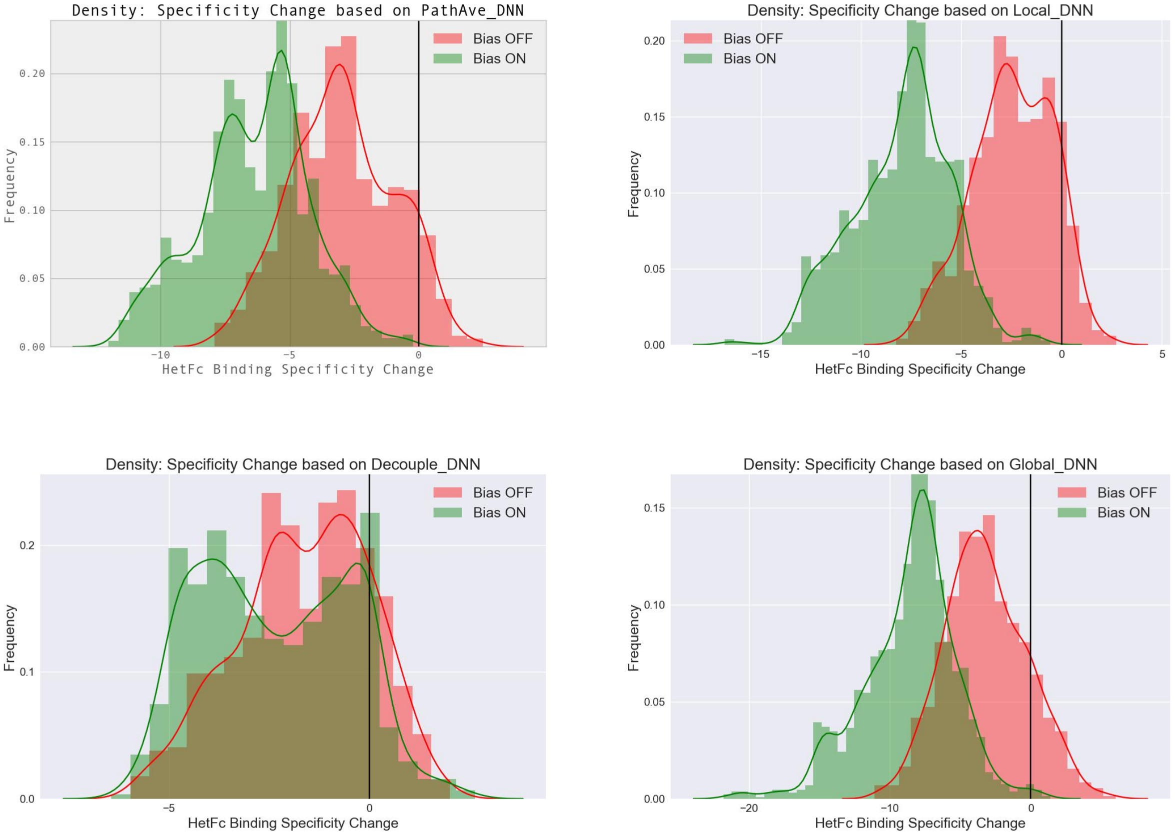

**FIGURE S10 "Average" (Ave) Specificity Metric change distributions from mutations generated by running automated sequence design on 74 target regions taken from our HetFc stability dataset.** Binding specificity metric changes due to mutation with respect to wild-type were evaluated from DNN-based energy functions. To evaluate the "PathAve_DNN" stability change metric for either a complex, ligand or receptor, 200 mutation paths were averaged over for each calculation of the stability change metric. Simulation protocols: wt_0.0_temp_1.0 (Bias OFF); wt_1.0_temp_0.25 (Bias ON).

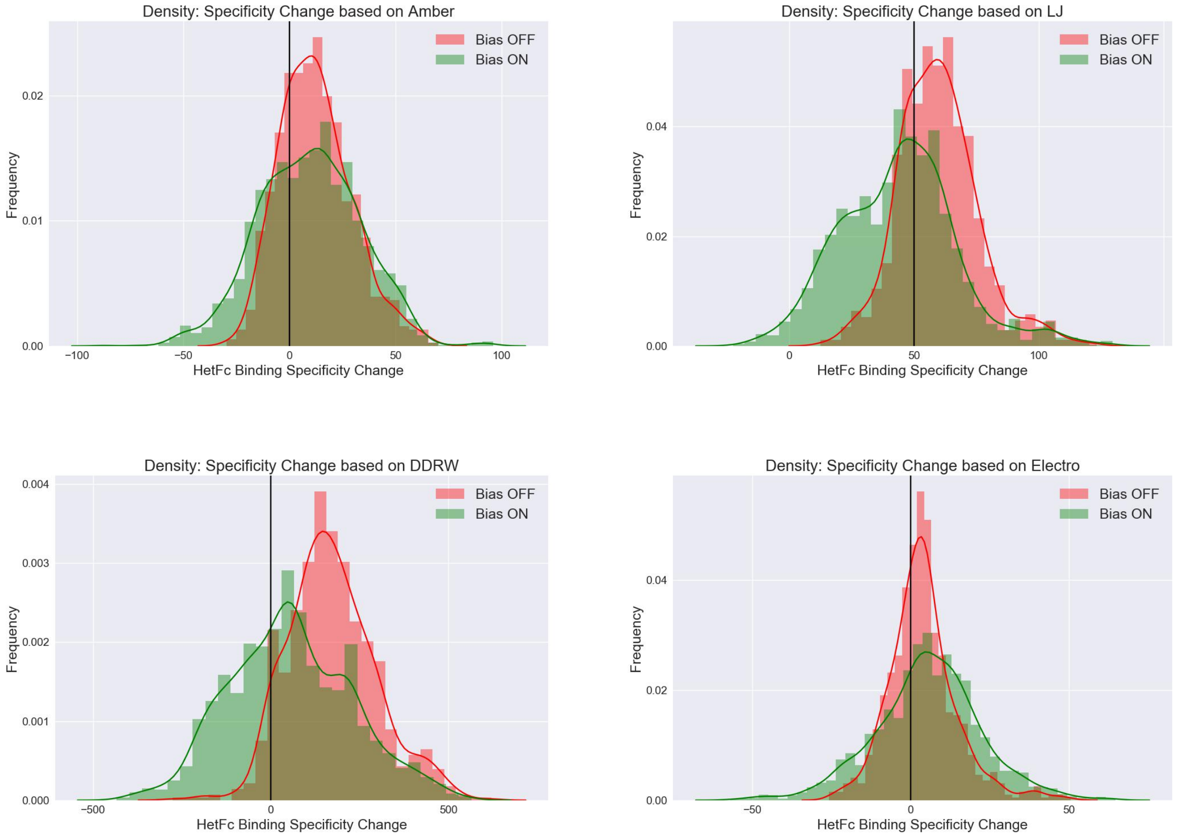

**FIGURE S11 "Minimum" (Min) Specificity Metric change distributions from mutations generated by running automated sequence design on 74 target regions taken from our HetFc stability dataset.** Binding specificity metric changes were calculated using physics-based and knowledge-based energy functions/stability metrics on structures output by our structural repacking workflow. Simulation protocols: wt_0.0_temp_1.0 (Bias OFF); wt_1.0_temp_0.25 (Bias ON).

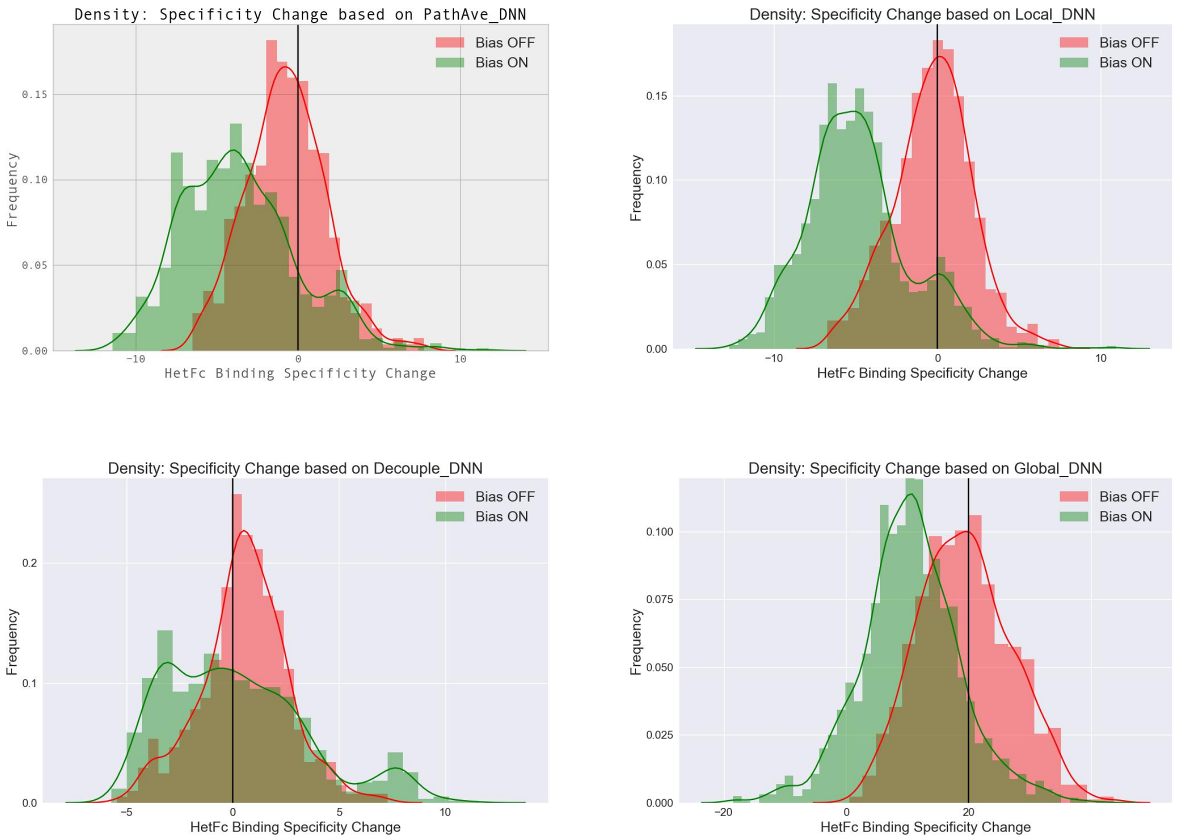

**FIGURE S12 "Minimum" (Min) Specificity Metric change distributions from mutations generated by running automated sequence design on 74 target regions taken from our HetFc stability dataset.** Binding specificity metrics were evaluated from DNN-based energy functions. To evaluate the "PathAve_DNN" stability change metric for either a complex, ligand or receptor, 200 mutation paths were averaged over for each calculation of the stability change metric. Simulation protocols: wt_0.0_temp_1.0 (Bias OFF); wt_1.0_temp_0.25 (Bias ON).

##### S.3.2 Automated HetFc Design on two example Target Regions, R1 and R2

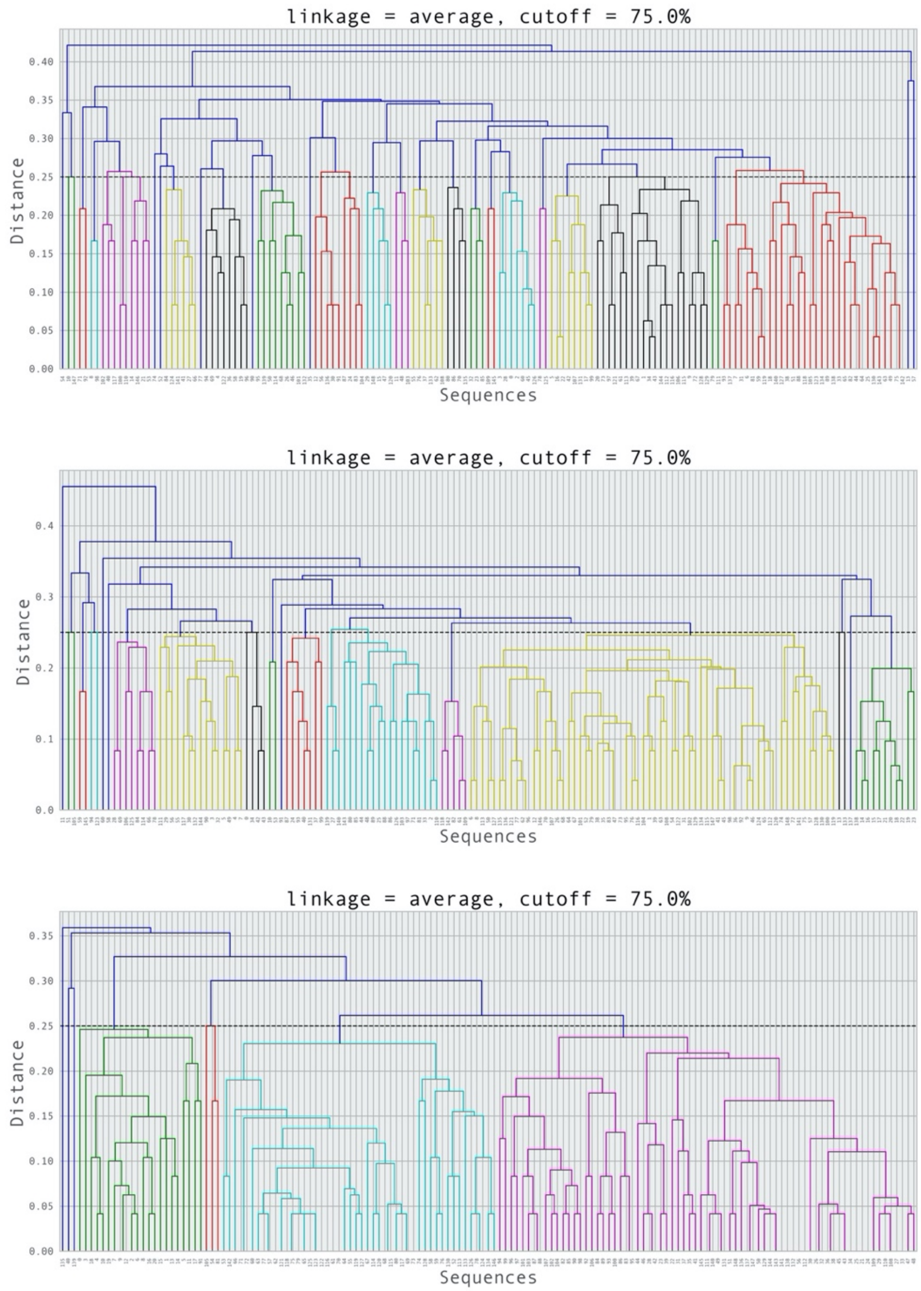

**FIGURE S13 Hierachical clustering of the simulation trajectory sequence snapshots generated by an automated specificity design run on the R1 target region of the Fc domain.** Each simulation run generated 150 simulation snapshots then a dendrogram was determined by hierarchical sequence clustering. Three simulation protocols were followed: (Top, bias OFF) wt_0.0_temp_1.0; (Middle, bias Moderate) wt_1.0_temp_1.0; and (Bottom, bias ON/strong) wt_1.0_temp_0.5. The sequence similarity cutoff was set to 75 percent: all sequences in a cluster (same colour *other than blue*) therefore were at least 75 percent sequence similar. Bottom panel: cyan branch, electrostatically driven design; purple branch, sterically driven (‘KiH’) design.

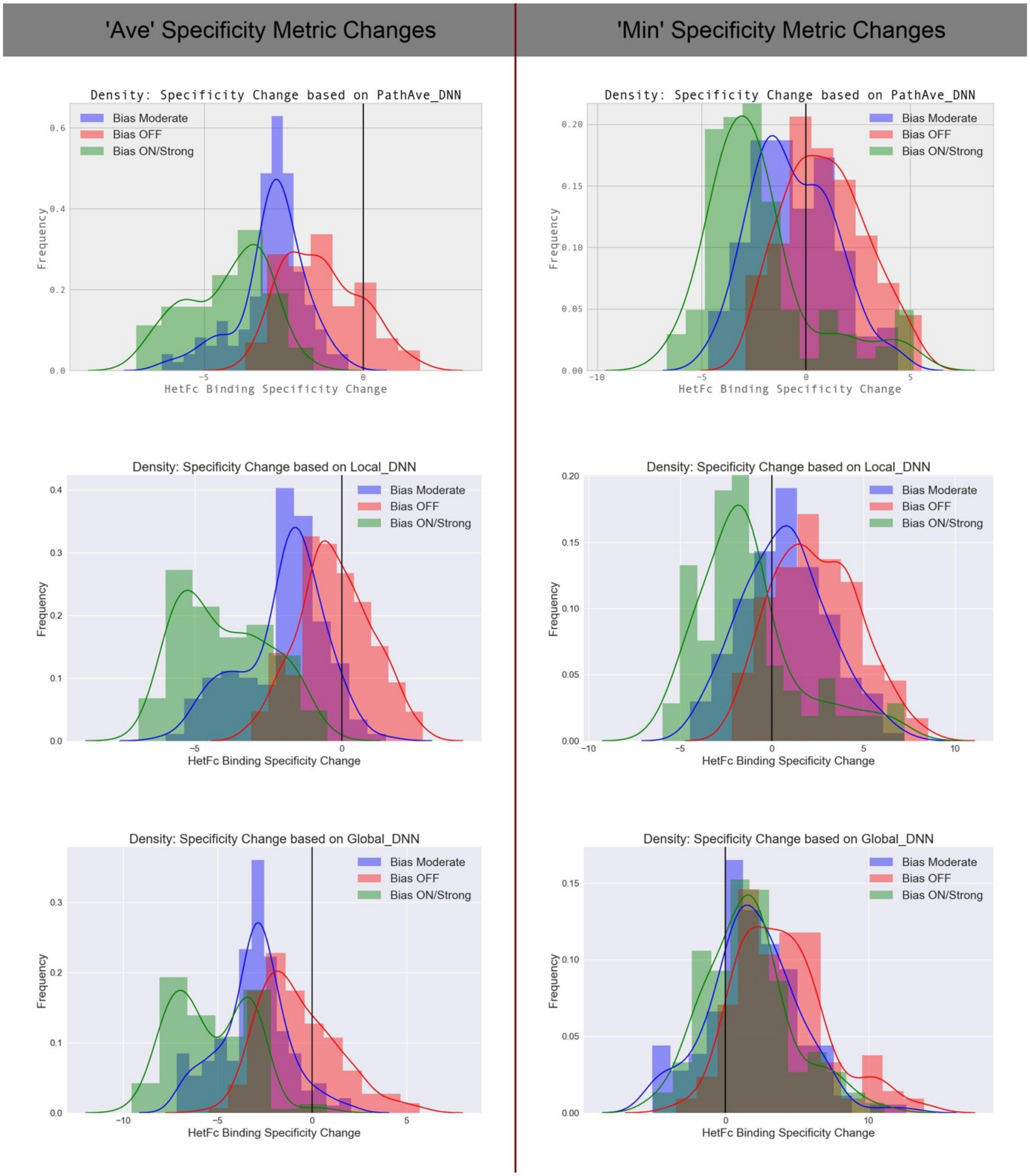

**FIGURE S14 Distributions of the change in the specificity metrics due to mutation relative to wild-type for sequences generated by an automated specificity design run on the R1 target region in the IgG1 Fc domain.**. Three simulation runs were performed each following a different protocol: (Red) wt_0.0_temp_1.0; (Blue) wt_1.0_temp_1.0; and (Green) wt_1.0_temp_0.5. Only unique $A_1B_2$ sequences / designs were used when determining a distribution: there were no repeated sequences. Specificity metrics were evaluated from DNN-based energy functions. To evaluate the “PathAve_DNN” stability change metric for either a complex, ligand, or receptor, 200 permutations to the order of the applied mutation swaps were averaged over for each calculation of the stability change metric.

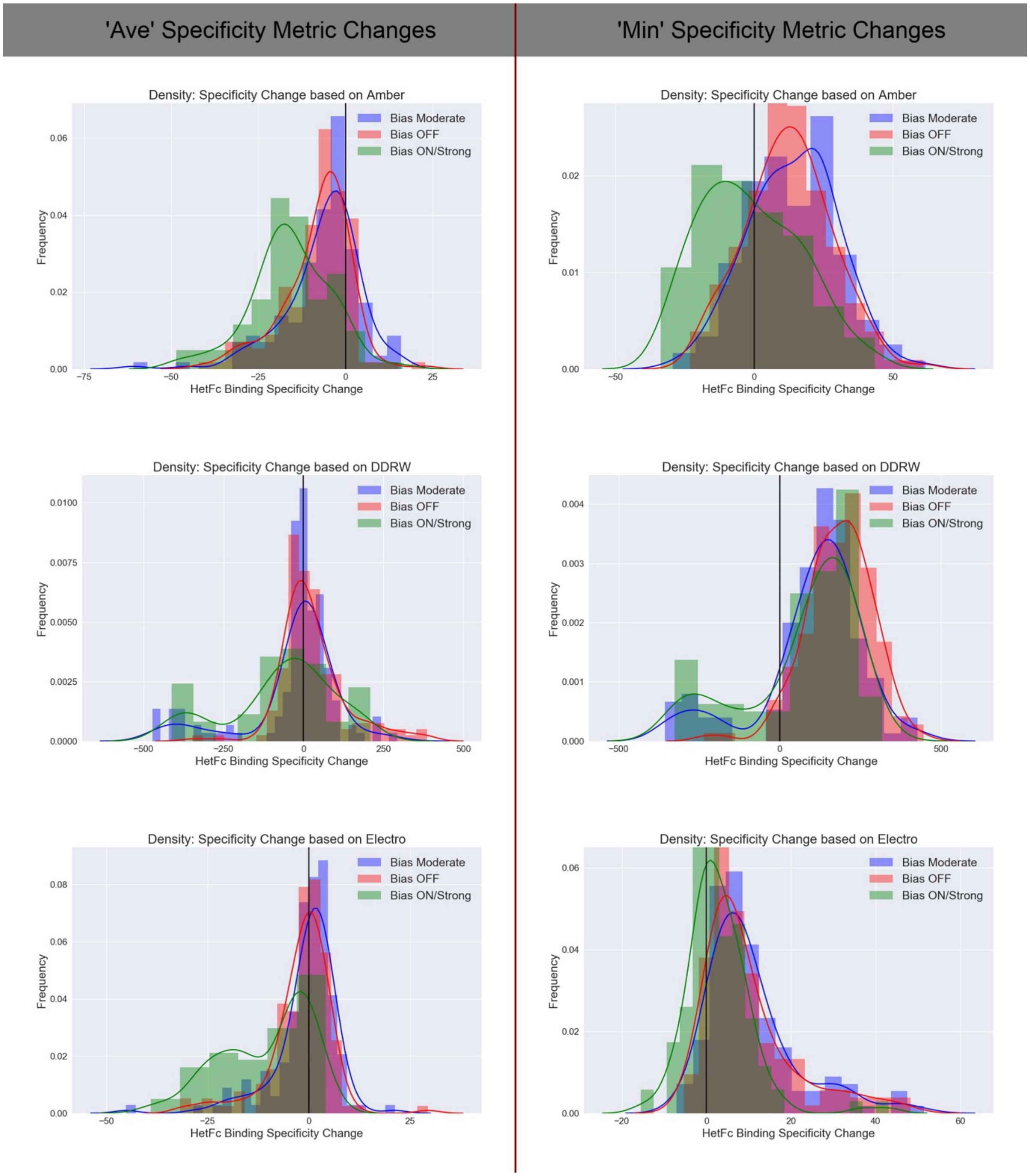

**FIGURE S15 Distributions of the change in the binding specificity metrics due to mutation relative to wild-type for sequences generated by an automated specificity design run on the R1 target region in the IgG1 Fc domain.** Specificity metric changes were calculated using physics-based and knowledge-based energy functions/stability metrics on structures output by our structural repacking workflow.

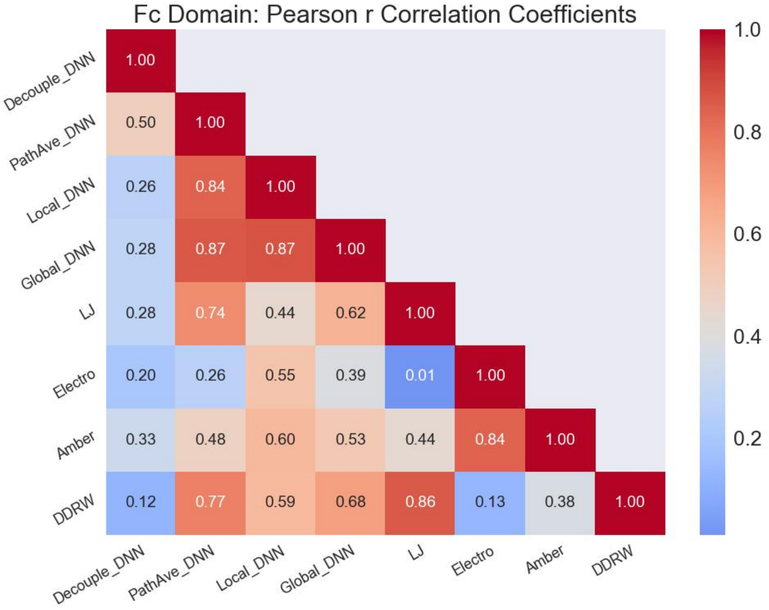

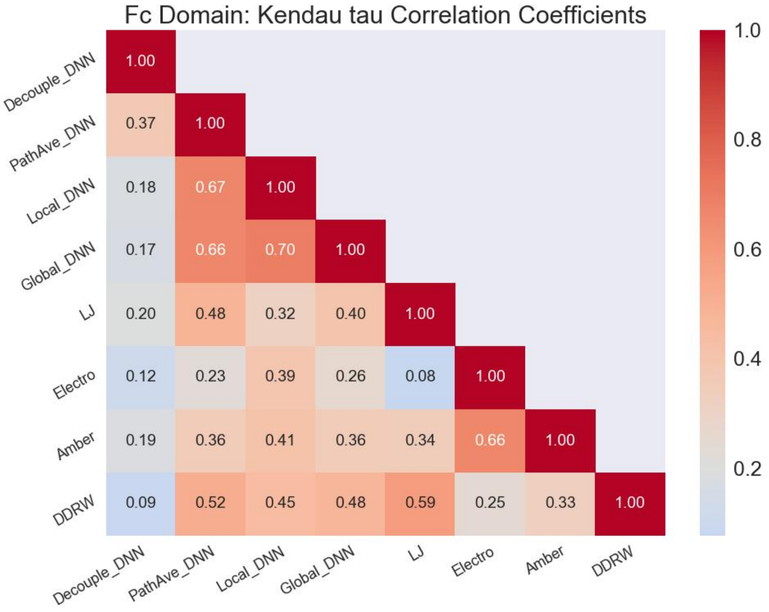

**FIGURE S16 Correlations (Pearson r and Kendall tau) between changes in "Ave" specificity metrics with respect to wild-type due to mutation for sequences generated by an automated specificity design run on the R1 target region in the Fc domain.** These changes were calculated using several different choices of underlying stability metric: standard physics-based and knowledge-based metrics (Amber, Electro, LJ, and DDRW) as well as DNN-based metrics (Local_DNN, Global_DNN, PathAve_DNN, and Decouple_DNN). All DNN-based metrics were evaluated using a DNN model, ZymeSwapNet, taking both backbone (BB) and side-chain (SC, neighbouring amino-acid identities) as features (ie, BB + SC; see discussions in Section 3). For the calculation of the "Decouple_DNN", swaps in any given mutation were treated as being decoupled: here when calculating the swap contribution to the stability change, residues neighbouring the swap site had their amino-acid identities set to wild-type. Simulation protocol is wt_1.0_temp_0.5.

| HetFc Design ( $A_1 B_2$ ) on Target Region R2 | Global_DNN $A_1 B_2$ | Global_DNN $A_1 B_1$ | Global_DNN $A_2 B_2$ |
|---|---|---|---|
| A/350T_A/351L_A/394T_A/395P_A/405F_A/407Y_B/350T_B/366T_B/392K_B/394T_B/397V_B/407Y | -14.644 | -14.644 | -14.644 |
| A/350V_A/351F_A/394S_A/395E_A/405S_A/407G_B/350M_B/366G_B/392L_B/394W_B/397S_B/407L | -16.625 | -12.876 | 23.847 |
| A/350V_A/351F_A/394T_A/395E_A/405S_A/407G_B/350I_B/366G_B/392L_B/394W_B/397R_B/407L | -14.714 | -12.644 | 26.303 |
| A/350V_A/351I_A/395G_A/405S_A/407C_B/350G_B/366L_B/392I_B/394T_B/397G_B/407C | -22.408 | -14.889 | -9.445 |

**TABLE S4 Absolute binding affinities for the Fc heterodimer and Fc homodimers.** A Knobs-into-Holes (KiH) and a disulphide HetFc selective binding sequences generated by automated specificity design run on the R2 target region and the R2 target region with position A/394 removed. Swaps to CYS and PRO were allowed. The wild-type reference design is colored in black and highlighted in bold. Simulation protocol was "wt_1.0_temp_0.25" (see discussion in Section 5.5). Notice, in the fourth HetFc sequence design(highlighted in brown), the presence of a CYS at symmetric positions A/407C and B/407C.

| HetFc Design ( $A_1 B_2$ ) on Target Region R2 | PathAve_DNN | Local_DNN | Global_DNN | Amber | DDRW |
|---|---|---|---|---|---|
| A/350V_A/351F_A/394S_A/395E_A/405S_A/407G_B/350M_B/366G_B/392L_B/394W_B/397S_B/407L | -4.732 | -6.029 | -3.749 | 15.813 | -210.99 |
| A/350V_A/351F_A/394T_A/395E_A/405S_A/407G_B/350I_B/366G_B/392L_B/394W_B/397R_B/407L | -3.613 | -4.076 | -2.070 | 16.853 | -312.97 |

**TABLE S5 "Min" Binding Specificity Metric change for the Fc heterodimer.** A Knobs-into-Holes (KiH) and a disulphide HetFc selective binding sequences generated by automated specificity design run on the R2 target region. Swaps to CYS and PRO were allowed. Simulation protocol was "wt_1.0_temp_0.25" (see discussion in Section 5.5).

Curiously, by dropping the A/394 from the R2 target region, such that this position had its amino acid identity fixed to wild-type THR, a separate automated specificity design placed a CYS at A/407 and another CYS at B/407, which were conserved along the simulation trajectory (Table S4). The proximity of these two sequence positions across the interface of the Fc domain strongly suggests that the pi-pi stacking A/407Y-B/407Y interaction was replaced by a disulphide bond, A/407C-B/407C. Our method does not explicitly predict disulphide bonds, but one can infer their presence via the strong probability of a CYS at two sufficiently close residue locations. However, given the symmetric positioning of these two CYS, it might be incorrect to say that the preferential heterodimer formation exhibited with this "HetFc design" was driven by a disulphide introduction: during binding it would require a disulphide forming for the $A_1B_2$ complex but not for the corresponding $A_1B_1$ and $A_2B_2$ complexes. We obtained some independent confirmation for the possibility of disulphide formation by running an internal tool with a custom energy function trained to detect pairs of residue positions where the insertion of a disulphide bond might be physically possible. In any case, this simulation result demonstrates the subtleties that could arise when enabling swaps to CYS.

### S.4 References

1. Xiang, Z. and Honig, B. Extending the Accuracy Limits of Prediction for Side-chain Conformations. *J. Mol. Biol.* 2001, 311, 421-430.
2. Besag, J.E. On the Statistical Analysis of Dirty Pictures. *J.R. Statist. Soc.* B **1986**, 48, 3, 259-302
